# Enabling tools for *Toxoplasma* glycobiology

**DOI:** 10.1101/401828

**Authors:** Elisabet Gas-Pascual, H. Travis Ichikawa, M. Osman Sheikh, M. Isabella Serji, Bowen Deng, Msano Mandalasi, Giulia Bandini, John Samuelson, Lance Wells, Christopher M. West

## Abstract

Infection by the protozoan parasite *Toxoplasma gondii* is a major health risk owing to its chronic nature, ability to reactivate to cause blindness and encephalitis, and high prevalence in human populations. Like nearly all eukaryotes, *Toxoplasma* glycosylates many of its proteins and lipids and assembles polysaccharides. Unlike most eukaryotes, *Toxoplasma* divides and differentiates in vacuoles within host cells. While their glycans resemble canonical models, they exhibit species-specific variations that have inhibited deeper investigations into their roles in parasite biology and virulence. The genome of *Toxoplasma* encodes a suite of likely glycogenes expected to assemble a range of *N*-glycans, *O*-glycans, a *C*-glycan, GPI-anchors, and polysaccharides, along with their requisite precursors and membrane transporters. To facilitate genetic approaches to investigate the roles of specific glycans, we mapped probable connections between 59 glycogenes, their enzyme products, and the glycans to which they contribute. We adapted a double-CRISPR/Cas9 strategy and a mass spectrometry-based glycomics workflow to test these relationships, and conducted infection studies in fibroblast monolayers to probe cellular functions. Through the validated disruption of 17 glycogenes, we also discovered novel Glc_0-2_-Man_6_-GlcNAc_2_-type N-glycans, GalNAc_2_- and Glc-Fuc-type O-glycans, and a nuclear O-Fuc type glycan. We describe the guide sequences, disruption constructs, and mutant strains, which are freely available to practitioners for application in the context of the relational map to investigate the roles of glycans in their favorite biological processes.

## INTRODUCTION

*Toxoplasma gondii* is a worldwide, obligate intracellular apicomplexan parasite that can infect most nucleated cells of warm-blooded animals (Kim and Weiss, 2004). Up to 80% of some human populations are seropositive (Pappas et al., 2009). Toxoplasmosis, the disease caused by *Toxoplasma*, is associated with encephalitis and blindness in individuals whose parasites are reactivated such as in AIDS and other immune-suppressed patients (Pereira-Chioccola et al., 2009). *In utero* infections can cause mental retardation, blindness, and death (Luft and Remington, 1992). *Toxoplasma* is transmitted by digesting parasites from feline feces (as oocysts) or undercooked meat (as tissue cysts). Once in the host, parasites convert to the tachyzoite form that disseminates to peripheral tissues (e.g. brain, retina, and muscle). The resulting immune response and/or drugs can control tachyzoite replication, but the parasite survives by converting into slow growing bradyzoites that encyst. Cysts sporadically burst, and the parasites convert to tachyzoites, whose unchecked growth results in cell and tissue damage (Lambert and Barragan, 2010; Blader et al., 2015). Currently, no *Toxoplasma* vaccine exists, anti-toxoplasmosis drugs have severe side effects, and resistance is developing to these drugs (Katlama et al., 1996; Dannamann et al., 1992; Baatz et al., 2006; Aspinall et al., 2002; Weiss and Kim, 2000). As infected individuals remain infected for life, new anti-*Toxoplasma* drugs are needed, and ideal targets are those present in the parasite but not in the host.

Apicomplexan proteins were for a time considered to be minimally glycosylated, but lectin-, metabolic labeling-, genomic- and mass spectrometry-based studies over the past three decades have refuted the original dogma in *Toxoplasma*, *Plasmodium* spp. (agent of malaria), *Cryptosporidium* spp. (agent of cryptosporidiosis), and others (Rodrigues et al., 2015; Bushkin et al., 2010; Heimburg-Molinaro et al., 2013; Macedo et al., 2010; Sanz et al., 2013). While less diverse than for most other eukaryotes, the major known glycan classes, including *N*-glycans, *O*-glycans, *C*-mannose, GPI-anchors, glycolipids, and polysaccharides, are represented in *Toxoplasma*. *N*-glycans, *O*-glycans, or GPI-anchors have been described on hundreds of *Toxoplasma* proteins (Luo et al., 2011; Fauquenoy et al., 2008; Wang et al., 2016; Bandini et al., 2016). Their functions have been little studied, but glycans have been implicated in invasion, O_2_-sensing, cyst wall formation, cyst persistence, and nutrient storage, each of which may be important for virulence (Luo et al., 2011; Debierre-Grockiego et al., 2010; Tomita et al., 2017; Rahman et al., 2016; Caffaro et al., 2013; Uboldi et al., 2015). Many glycosyltransferase genes have low fitness scores when disrupted in a genome-wide survey (Sidik et al., 2016), indicating unknown but vital biological functions for the parasite (Fig. 1).

**Fig. 1.**
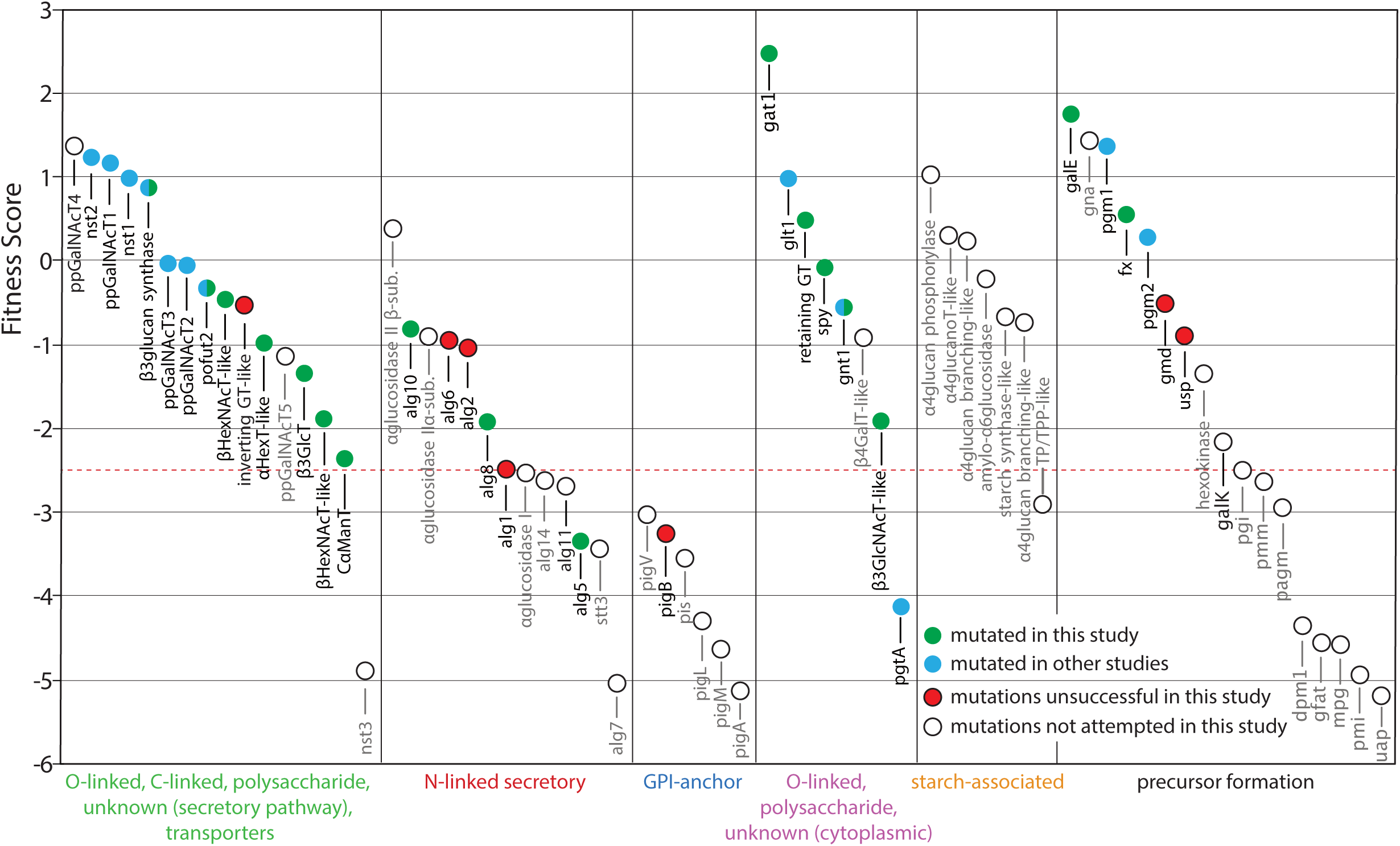
Fitness of glycogenes. Fitness data for glycogenes for invasion of and growth in human fibroblasts were extracted from Sidik et al. (2016) and graphed according to rank order of fitness value for separate categories of glycosylation as indicated at the bottom. Fitness scores are based on a log2 scale and values below -2.5 (red dashed line) have increasing likelihood of being essential. Filled data points represent genes for which mutants are available: green-from this study, blue-from other studies. Red fill refers to unsuccessful attempts from this study, and unfilled circles indicate no mutants attempted (but dual guide plasmids, or pDGs, available, in black) or known (in gray). Genes names correspond to listings in Table 1.

Glycosylation associated drugs are beginning to make an impact in human health, and there is reason to anticipate success in combating pathogens with their distinctive glycan synthetic pathways. Unique glycoconjugates are prime candidates for vaccine development and as markers for diagnostic monitoring. But before the full potential of these strategies can be realized, more structure-function studies are needed to guide selection of the most promising targets. Progress in exploiting these strategies has been inhibited by the uniqueness of the glycomes of *Toxoplasma* and other parasites, and lack of information about the genetic basis of its non-template driven mechanisms of assembly.

Further understanding of the roles of glycans will benefit from genetic approaches to manipulate their biosynthesis. Sufficient knowledge currently exists to allow informed predictions of glycogenes associated with glycan assembly, and mass spectrometry-based technologies for interpreting glycomic consequences of glycogene perturbations are available. This report describes the generation of a collection of CRISPR/Cas9-associated guide DNAs that are suitable for disrupting glycogenes, using an approach that is expected to be suitable for strains regardless of their NHEJ (Ku80) status, and presents the results of trials on the type 1 RH strain. A streamlined mass spectrometry workflow was implemented to benchmark the new studies, which led to some revisions of existing models and new structures including novel N-glycans, a novel mucin-type O-glycan, and a novel nuclear O-Fuc. Finally, the glycogene mutants were screened in a cell culture growth assay for information about their physiological significance. It is expected that the availability of these findings and associated resources will help stimulate interrogation of the role of glycosylation in future functional studies.

## EXPERIMENTAL PROCEDURES

### Glycogene prediction

To search for candidate glycosyltransferases (GTs) that contribute to the *Toxoplasma* glycome, we expanded approaches previously implemented in *Dictyostelium discoideum* (West et al., 2005) and *D. purpureum* (Sucgang et al., 2011). The predicted proteome (8460 proteins) from Toxodb.org was analyzed by i) the SUPERFAMILY server, which assigns protein domains at the SCOP ‘superfamily’ level using hidden Markov models; ii) dbCAN, an automated <u>C</u>arbohydrate-active enzyme <u>AN</u>notation database which utilizes a CAZyme signature domain-based annotation based on a CDD (conserved domain database) search, literature curation, and a hidden Markov model; iii) the Pfam database; and iv) Bernard Henrissat (Marseille) who supervises the CAZy database (Lombard et al., 2014). This yielded 39 GT-like coding regions. No gene candidates for DNA glucosylation as characterized in trypanosomatids (Bullard et al., 2014) are apparent in apicomplexans. The search was expanded to include genes predicted to contribute to the synthesis of sugar nucleotide precursors and their transport into the secretory pathway, genes that contribute to *N*-glycan processing, and genes associated with processing of starch, yielding 28 additional glycogenes.

### Parasites and cell culture

Human foreskin fibroblasts (HFF, ATCC SCRC-1041) and HFF immortalized with human telomerase reverse transcriptase (hTERT, ATCC CRL-4001) were maintained in Complete Medium, which consisted of Dulbecco’s Modified Eagle Medium (DMEM, Corning Inc., 15-1013-CV) supplemented with 10% (v/v) fetal bovine serum (FBS, Corning, 35-015-CV), 2 mM L-glutamine (Corning, 25-005-CI), and 100 IU/ml penicillin/100 µg/ml streptomycin (Corning, 30-002-CI). *Toxoplasma gondii* strains RH and Pru, and their derivative strains, were routinely maintained on hTERT cells in Complete Medium containing 1% (v/v) FBS in a humidified incubator with 5% CO_2_ at 37°C.

Intracellular parasites were prepared by forcing well-infected fibroblast cells through a 27-gauge needle attached to 10 ml syringe 5 times for hTERT cells and 3 times for HFF cells and collected by centrifugation at 2000 × *g* for 8 min at 4°C.

Parasite freezer stocks were prepared from a T25 flask culture of hTERT or HFF monolayer that were 30 to 50% lysed. Scraped cells were collected and pelleted as described above and resuspended in 1-1.5 ml ice-cold DMEM:FBS (1:1). After incubating 10 min on ice, an equal volume of ice-cold DMEM:DMSO (Sigma, D2650) in a 4:1 ratio was added dropwise to the cell suspension. Cell suspensions were slowly frozen in 1-ml cryo-vials at -80°C and transferred to liquid N_2_.

### Dual-guide plasmid construction and DHFR cassette amplification

Plasmid 2 and Plasmid 3 were derived from pU6-Universal (Sidik et al., 2016; Addgene plasmid #52694) by the introduction of a novel XhoI site either upstream of the U6 promoter, or between the guide RNA scaffold and the NsiI site, respectively. Plasmid 2 (p2) was constructed by replacing the PvuI-PciI fragment of pU6-Universal with a new PvuI-PciI fragment generated by PCR of pU6-Universal with Plasmid 2 FOR and Plasmid 2 REV (which contains an XhoI site) (Table S3), using pU6-universal as the template. Similarly, Plasmid 3 (p3) was constructed by replacing the HindIII-NsiI fragment with the HindIII-NsiI fragment of a PCR amplicon generated using Plasmid 3 FOR and Plasmid 3 XhoI (which contains an XhoI site). All enzymatic manipulations of plasmid DNA were performed as described by the manufacturer, and all PCR amplification reactions were performed with Q5 DNA polymerase (NEB, M0491L). p2 and p3 were prepared in *dam*^*–*^/dcm^*–*^ Competent *E. coli* (NEB, C2925I). Guide sequences were selected using the Eukaryotic Pathogen CRISPR guide RNA/DNA Design Tool (EuPaGDT; Peng and Tarleton, 2015; http://grna.ctegd.uga.edu) based on the calculated efficiency score against GT1 genome (and/ or Me49 genome) in the latest available version of ToxoDB (http://toxodb.org/toxo/). Guide sequences for each gene were cloned into p2 and p3, as described in Addgene protocol with minor modifications. Briefly, annealed synthetic “top” and “bottom” ssDNA guide oligonucleotides flanked by BsaI compatible 4-nt overhangs were directionally cloned and insertion was confirmed by PCR reactions using the “top” strand guide oligonucleotide and a downstream primer, Plasmid 1 REV. Dual-guide plasmids (pDG) were constructed by transferring the XhoI-NsiI fragment of p2 into p3 that was double digested with XhoI and NsiI (Fig. 2). Plasmids were prepared in transformed Top10 cells (Thermo Fisher Scientific) using a ZR Plasmid Miniprep-Classic (Zymo Research, D4016) kit. Finally, the inserted guide sequences were confirmed by sequencing the pDG plasmid with primers gRNA For and gRNA Rev (Table S3). All available plasmids are referred to in Table S1 and defined in Table S4.

**Fig. 2.**
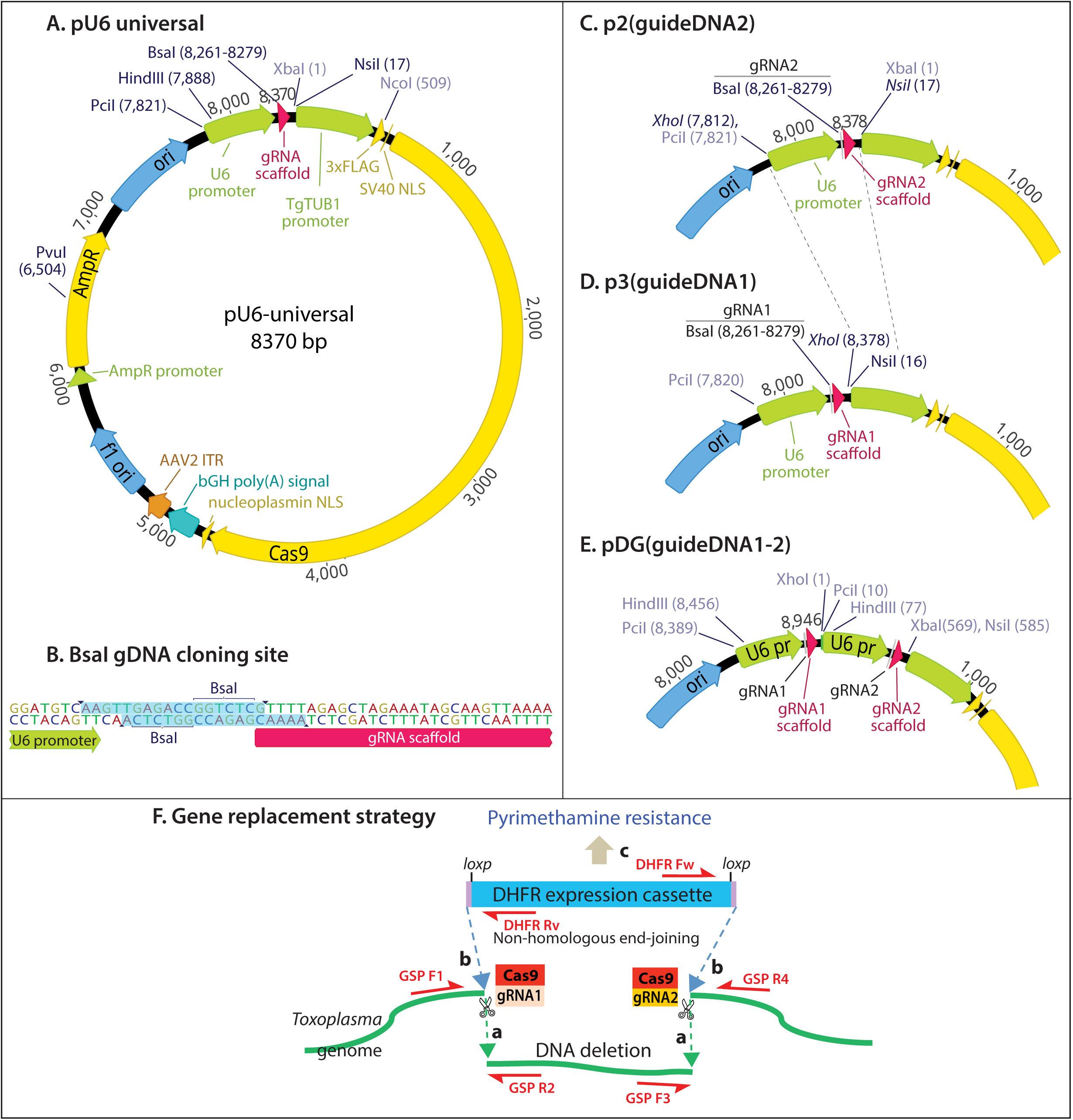
Plasmids and gene replacement. The previously described pU6-Universal plasmid **(A)**, designed for single CRISPR/Cas9 editing, contains a BsaI site for convenient introduction of a synthetic duplex guide DNA **(B)**. pU6 was modified by the addition of a XhoI site at the 5’-end of its guide DNA cassette to generate p2 **(C)**, or its 3’end to generate p3 **(D)**. Thus the guide2 cassette of p2 could be conveniently excised from and re-ligated into p3 to generate the dual guide plasmid pDG **(E)**. **(F)** Gene replacement strategy. Co-transient transfection of pDG, expressing the gRNAs and Cas9, with the DHFR amplicon, results in the excision of the intervening genomic DNA (a) and ligation of the DHFR amplicon by non-homologous end joining (b). The desired replacement clones are selected for in the presence of pyrimethamine (c) and screened by PCR using gene specific primers (GSP) flanking the excision sites.

The DHFR expression cassette amplicon was prepared using primers (DHFR-Fw or DHFR-For LoxP2 and DHFR-Rv) (Table S3) and purified using a DNA Clean & Concentrator-5 kit (Zymo Research, D4003). For transfection, pDG and the DHFR expression amplicon were EtOH-precipitated, rinsed with 70% EtOH, and resuspended in Cytomix Buffer (120 mM KCl, 0.15 mM CaCl_2_, 10 mM K_2_HPO_4_/KH_2_PO_4_, 25 mM HEPES-NaOH, 2 mM EDTA, 5 mM MgCl_2_, final pH 7.6).

### Transfection and cloning

Transfection was performed as described (Striepen and Soldati, 2007). Briefly, 10 million freshly prepared intracellular parasites were electroporated with 1 µg DHFR expression amplicon ± 10 µg pDG, or no DNA, in Cytomix Buffer plus 100 mM each of ATP and glutathione in a final vol of 400 µl in a 2 mm-wide cuvette (Bulldog BIO, 12358-346), at a setting of 1.5 kV, 25 µF in a Gene Pulser Xcell (Bio-Rad). After 10 min of incubation at room temperature, treated parasites were diluted into 5 ml of Complete Medium with 10% FBS and inoculated onto confluent HFFs in a T25 flask. After standard incubation overnight, the medium was replaced with 5 ml of Complete Medium with 10% FBS containing 1 µM pyrimethamine (Sigma, 46703) for selection. If the majority of HFF cells started lysing before the mock-transfected parasites died, the parasites were transferred to another T25 flask with hTERT monolayer for further selection in Complete Medium with 1% FBS containing 1 µM pyrimethamine until no more visibly viable mock-transfected parasites were observed.

Pyrimethamine-resistant clones were isolated by limiting dilution on confluent HFF in Complete Medium with 1% FBS containing 1 µM pyrimethamine. Six serial two-fold dilutions starting with 25 parasites per well were prepared in octuplet in a 96-well plate and grown for 6 to 10 d. Single clones were expanded in hTERT monolayers in T25 flasks for genomic DNA isolation and freezer stock preparation.

### Analysis of *Toxoplasma gondii* glycogene loci

Genomic DNA was prepared from extracellular parasites using a Quick DNA Miniprep Plus kit (Zymo Research, D4069), eluted in 100 µl of DNA Elution Buffer and stored at 4°C in the DNA Elution Buffer. Q5 DNA polymerase was used to PCR amplify 10-20 ng of genomic DNA with primer sets flanking expected double stranded break sites (Table S2) and near the ends of DHFR amplicon (Table S3) to characterize the glycogene loci disruption.

### Plaque assay

Plaque assays were performed on 100% confluent, tightly packed HFF monolayers under 3 ml Complete Medium, 1% FBS without pyrimethamine in a 6-well plate. Freshly lysed-out intracellular parasites were inoculated at 100 cells/well, and the medium was replaced after 3 h by aspiration. On d 6, cultures were aspirated and rinsed 3× with 3 ml PBS (Corning, 21-040-CV) before fixation with -20°C 100% methanol for 5 min followed by air drying. The monolayer was stained with 1 ml 2% (w/v) crystal violet, 0.8% ammonium oxalate (w/v) in 25% ethanol for 5 min at room temperature, rinsed 3× with PBS, and air dried. Mutant and control (RH) strains were always compared on the same plate. Monolayers were scanned by Odyssey CLx Imaging System (Li-Cor) and plaque areas were measured using ImageJ software (National Institute of Health, USA). Mutant plaque areas were referenced to control plaque areas from the same plate, and all control plaque areas were normalized to a value of one for presentation. Data were presented and statistically analyzed using GraphPad Prism version 6.

### Preparation of parasites

The type 1 RH strain (w/t) and mutant strains were grown on confluent hTERTs in Complete Medium supplemented with 1% FBS. The medium was collected after 36-48 h, and spontaneously lysed-out tachyzoites were harvested by centrifugation (8 min, 2000 × *g*, room temperature). The pellets were syringe-lysed (through a 27G needle) three times, filtered through a 3 µm Nuclepore^®^ filter (Whatman) followed by 2 rinses of DMEM to remove host cell debris. Pellets were resuspended and centrifuged 3× with ice-cold PBS, frozen in liquid N_2_, and stored at -80°C. 125 mg wet weight of cells (∼5 × 10^9^ cells) were typically recovered from 14 T175 flasks.

### Preparation of delipidated total cell pellets and Folch-partition

Cell pellets (∼5 × 10^9^ cells) were delipidated, and total protein powder was prepared as described (Aoki et al., 2007). Briefly, cell pellets were disrupted by Dounce homogenization in a mixture of ice-cold MeOH and H_2_O. The resulting suspension was transferred to a screw-top 8-ml glass tube and adjusted with water and CHCl_3_ to a final ratio of CHCl_3_:MeOH:H_2_O (4:8:3;v:v:v). The extract was incubated for 3 h at room temperature with end-over-end agitation. The insoluble proteinaceous material was collected by centrifugation and re-extracted three times in CHCl_3_/MeOH (2:1;v:v) and twice in CHCl_3_:MeOH:H_2_O (4:8:3;v:v:v). The final pellet was dried under a stream of N_2_, washed with acetone, dried, weighed (typically ∼25-30 mg), and stored at -20°C until analysis. The clarified lipid extract was subjected to Folch-partitioning (Folch et al., 1957). The resulting aqueous fraction (GIPLs) and organic fraction (other lipids) were dried under a stream of nitrogen and stored separately at -20°C.

### Enrichment in GPI-anchored proteins and GPI-glycan core release

GPI-anchored proteins were enriched from delipidated pellets by partitioning in butan-1-ol (n-BuOH, 99%; Alfa Aesar, Tewksbury, MA) as described (Almeida et al., 1994; Koeller et al., 2014). Briefly, 20 mg of protein powder was extracted 3× with 4 ml H_2_O saturated with n-BuOH (∼9%) for 4 h at room temperature. The n-BuOH-saturated water fractions were pooled and extracted with an equal vol of n-BuOH. After centrifugation for 15 min at 3500 × *g* at room temperature, upper butanolic and lower aqueous phases were separately dried and stored at -20°C. Butanolic fractions, enriched in GIPLs, were saved for future analysis. Aqueous fractions, containing GPI-anchored proteins, were treated with 100 µl 0.5 M NH_4_OH (in H_2_O) for 6 h at 4°C to release fatty acids. The sample was then dried under a N_2_ stream, redissolved in H_2_O, and dried again twice. The glycan core was released by treatment with 50 µl 48% aqueous HF at 4°C for 12 h (Masuishi et al., 2016), resuspended in 500 µl 0.05% trifluoroacetic acid (ThermoFisher Scientific, 28904) in H_2_O, filtered through a C_18_-SepPak to capture protein and lipid, and dried under vacuum. The sample was subjected to N-acetylation by addition of 10 µl of pyridine (Sigma, 360570) and 50 µl of acetic anhydride (Sigma, 539996) in 0.5 ml of MeOH, incubation for 30 min at room temperature (Kozulic et al., 1979), and drying under a stream of N_2_.

### Preparation of *N*-linked glycans

*N*-linked glycans were prepared as described (Aoki et al., 2007). 2 to 5 mg of protein powder was resuspended in 450 µl of trypsin digestion buffer (20 mM NH_4_HCO_3_, natural pH, 1 M urea) with brief bath sonication. 25 µl each of 20 ng/µl trypsin (Sigma, T8003) and 10 µg/µl chymotrypsin (Sigma, C4129) was added at a final ratio ∼1:20 protease:protein and incubated for 18 h at 37°C. The reaction was boiled for 5 min, cooled and centrifuged to remove insoluble material. The supernatant was adjusted to 5% (v/v) acetic acid (HAc) and loaded onto a Sep-Pak C_18_ cartridge (Sep-Pak^®^ Vac 1cc C_18_ cartridge, Waters, Milford, MA), that had been pre-equilibrated by washing with >3 ml of 100% acetonitrile and >3 ml of 5% HAc, and washed with 10 column vol of 5% HAc. Glycopeptides were eluted stepwise, as described (Alvarez-Manilla et al., 2010), with 2 vol 20% 2-propanol in 5% HAc, 2 vol 40% 2-propanol in 5% HAc and 3 vol 100% 2-propanol. The 2-propanol fractions were pooled, evaporated to dryness by vacuum centrifugation, resuspended in 50 µl of 20 mM Na phosphate buffer (pH 7.5), and digested with PNGaseF (NEB, P0705S) for 18 h at 37°C. Reaction mixtures were evaporated to dryness, resuspended in 500 µl 5% HAc and loaded onto a Sep-Pak C_18_ cartridge column. The column run-through and an additional wash with 3 column vols of 5% HAc, containing released oligosaccharides, were collected together and evaporated to dryness. Core α3-linked Fuc, which interferes with PNGaseF action, has not been detected on *Toxoplasma N*-glycans consistent the absence of candidate α3FucT encoding genes in the genome.

For samples to be treated with α-mannosidase, the PNGaseF digestion was performed instead in 25 mM NH_4_HCO_3_ (pH 7.8) and taken to dryness. The sample was resuspended in either 50 µl of ∼8 U/ml of Jack Bean α-mannosidase (Prozyme, GKX-5010) in 20 mM NaAc (pH 5.0), 0.4 mM Zn^2+^, or of ∼5 mU/ml of *Aspergillus satoi* α-mannosidase (Prozyme, GKX-5009, in 100 mM NaAc (pH 5.0), and digested for 18 h at 37°C. The reaction mixture was evaporated to dryness, resuspended in 500 µl 5% HAc, purified through a Sep-Pak C_18_ cartridge, and again evaporated to dryness.

### Preparation of *O*-linked glycans

*O*-linked glycans were obtained by reductive β-elimination as described previously (Aoki et al., 2008) with minor modifications. Briefly, ∼5 mg of protein powder was resuspended with bath sonication in 500 µl of 50 mM NaOH (diluted from 50% stock; Fisher Scientific, SS254-500) containing 1 M NaBH_4_ (Sigma, 213462-25G), and incubated for 18 h at 45°C in an 8-ml glass tube sealed with a Teflon-lined screw top. The reaction mixture was then neutralized with 10% HAc and desalted by applying to a 1-ml bed of Dowex 50W-X8 (H^+^ form; Sigma, 217506) and washing with 5 bed volumes of 5% HAc. The flow-through fraction, containing released oligosaccharide alditols, was evaporated to dryness. Borate was removed as an azeotrope with MeOH by adding 0.5 ml of 10% HAc in MeOH, drying under a N_2_ stream at 42°C, and repeating three additional times. To remove residual peptide and reagent contaminants, the dried material was resuspended in 500 µl of 5% HAc and loaded onto a C_18_-SepPak as above. Released oligosaccharide alditols were recovered by collecting the column run-through and an additional 2 ml of wash with 5% HAc. The run-through and wash were combined and evaporated to dryness.

### Mass spectrometry of permethylated glycans – direct infusion

The released glycans from the original 5-20 mg of protein powder were permethylated, as described (Anumula and Taylor, 1992), and the final CH_2_Cl_2_ phase was dried down under N_2_ and redissolved in 100 µl of MeOH. 10 µl of the glycan sample was combined with 35 µl of MS Buffer (1 mM NaOH in 50% MeOH) and 5 µl of ^13^C-permethylated isomaltopentaose (DP5), as an internal standard at a final concentration of 0.2 pmol/µl. Samples were directly infused (1 µl/min) into an Orbitrap mass spectrometer (LTQ Orbitrap XL or Orbitrap Elite, ThermoFisher Scientific) using nanospray ionization in positive ion mode, with the capillary temperature set to 200°C. The Full FTMS was collected and the Top 10 most intense peaks were selected for MS^2^ fragmentation using collision-induced dissociation (CID) at 40% collision energy. Once an ion was fragmented, the *m*/*z* was automatically put on an ‘exclusion list’ to prevent re-analysis in the run. MS^2^ product fragments matching a preselected list (Table S5) were selected for MS^3^ CID fragmentation at the same collision energy.

### Mass spectrometry of permethylated glycans – nLC-MS/MS

10 µL of permethylated glycans from above were combined with 2 µL of an internal standard mix [^13^C-permethylated isomaltose series (DP4, DP5, DP6, and DP7) at a final concentration of 0.2 pmol/µl each] and 38 µL of LC-MS Buffer A (1 mM LiAc, 0.02% HAc). 5 µl was injected into a PepMap Acclaim analytical C18 column (75 µm × 15 cm, 2 µm pore size) maintained at 60°C in an Ultimate 3000 RSLC (ThermoFisher Scientific/Dionex), which was coupled to a ThermoScientific Velos Pro Dual-Pressure Linear Ion Trap mass spectrometer. For *N*-Glycans, after equilibrating the column in 99% LC-MS Buffer A and 1.5-min ramp up to 45% LC-MS Buffer B [80% (v/v) acetonitrile, 1 mM LiAc, 0.02% HAc], separation was achieved using a linear gradient from 45% to 70% Buffer B over 150 min at a flow rate of 300 nl/min. For *O*-Glycans, after equilibrating the column in 99% LC-MS Buffer A and a 1.5-min ramp to 30% Buffer B, separation was achieved using a linear gradient from 30% to 70% Buffer B over 150 mins at a flow rate of 300 nl/min. The column was regenerated after each run by ramping to 99% Buffer B over 6 min and maintaining at 99% Buffer B for 14 min, before a 1-min ramp back down to 99% Buffer A. The effluent was introduced into the mass spectrometer by nanospray ionization in positive ion mode via a stainless-steel emitter with spray voltage set to 1.8 kV and capillary temperature set at 210°C. The MS method consisted of first collecting a Full ITMS (MS^1^) survey scan, followed by MS^2^ fragmentation of the Top 3 most intense peaks using CID at 42% collision energy and an isolation window of 2 *m*/*z*. Dynamic exclusion parameters were set to exclude ions for fragmentation for 15 s if they were detected and fragmented 5 times within 15 s.

### Glycan annotation

Structural assignments for the glycans detected at the reported *m*/*z* values were based on the compositions predicted by the exact mass (±0.1 amu) of the intact molecule, the presence of diagnostic MS^2^ and MS^3^ fragment ions that report specific glycan features, and the limitations imposed on structural diversity by known glycan biosynthetic pathways. The GRITS toolbox and its GELATO annotation tool (Ranzinger et al., 2017) were used to help identify glycan compositions using customized glycan databases and with the following tolerance settings: MS^2^ accuracy 500 ppm, max number of adducts (charge states) = 3.

### Western blotting and immunofluorescence analysis

Whole cell pellets (3 × 10^6^ cells/lane) were solubilized in Laemmli sample buffer containing 50 mM dithiothreitol, subjected to SDS-PAGE on 4–12% preformed gels in MES buffer (NuPAGE Novex, Invitrogen), and Western blotted onto nitrocellulose membranes using an iBlot system (Invitrogen). Blots were blocked with 5 mg/ml bovine serum albumin in PBS. For Tn antigen, blots were probed with anti-Tn mAbs 1E3 and 5F4 (Steentoft et al., 2011), which were validated for specificity by comparison of reactivity with wild-type and β3GalT (Core 1)-KO mouse colon samples generously provided by Lijun Xia (OMRF), and a 1:10,000-fold dilution of Alexa-680-labeled rabbit anti-mouse IgG secondary antibody (Invitrogen). For *O*-Fuc, blots were probed with biotinylated *Aleuria aurantia* lectin (AAL) and ExtrAvidin HRP (Sigma) and imaged on an ImageQuant LAS4000 imager (GE Healthcare) as previously described (Bandini et al., 2016). Blots were imaged on a Li-Cor Odyssey infrared scanner and processed in Photoshop, with contrast maintained at γ=1.

Parasites were analyzed by immunofluorescence with AAL as previously described (Bandini et al., 2016).

## RESULTS AND DICUSSION

### Predicted glycogenes

Our algorithms predicted 38 GT-like genes expected to contribute directly to the assembly of *N*-, *O*- and *C*-glycans on proteins, GPI anchors and other glycolipids, starch, and a wall polysaccharide. An additional five genes are likely to be involved with starch assembly and disassembly, one in GPI processing, three in *N*-glycan processing, 16 in the assembly of sugar nucleotide precursors from simple sugars, and three in transport of sugar nucleotides into the secretory pathway. Table 1 summarizes characteristics of the candidates, organized according to the class of glycoconjugates for which they are expected to contribute, along with findings from the literature and studies described below that provide support for the predicted functions. All but eight of the genes are correlated graphically with known or predicted glycosylation pathways in Figs. 4-8.

**Table 1.**
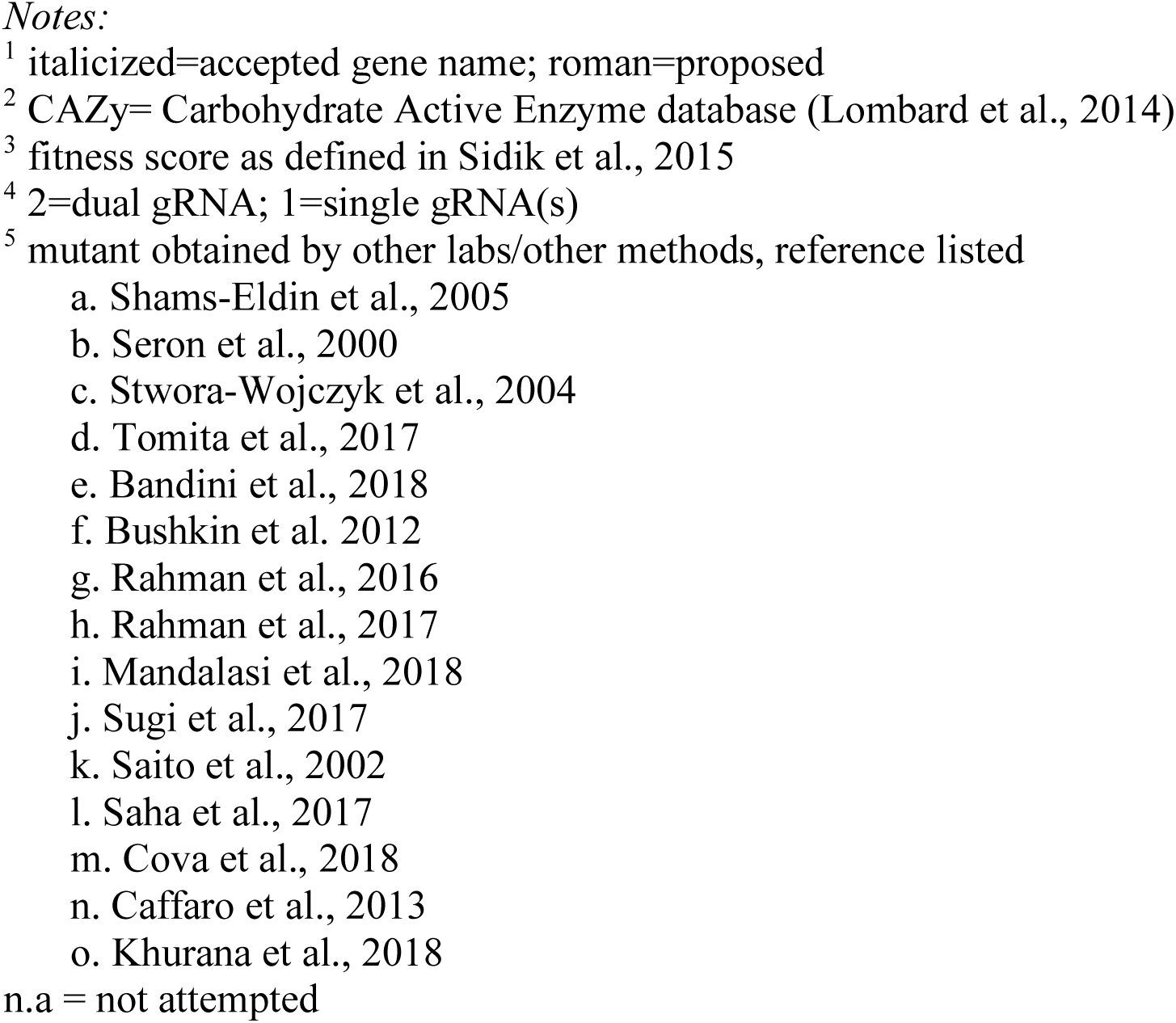
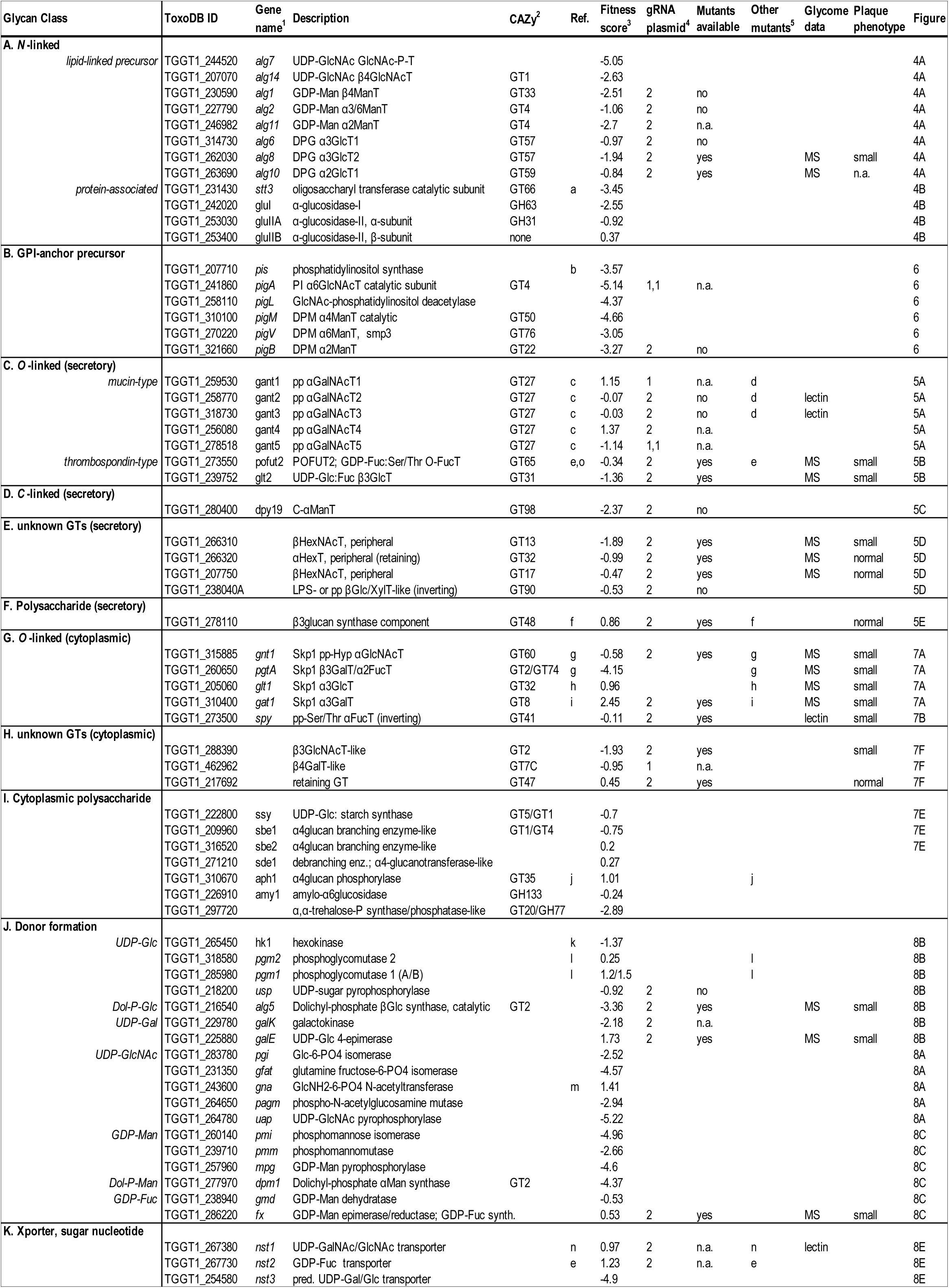
*Toxoplasma* glycogenes and their characteristics. Summary of documented and predicted genes contributing directly and indirectly to the parasite glycome. Genes are organized according to glycan type to which they are expected to contribute. Characteristics and evidence regarding their functions from this project and others are summarized.

For an initial indication of the cellular roles of the glycogenes, we analyzed data from a published genome-wide CRISPR gRNA-based screen for genes that contribute to fitness in a fibroblast monolayer infection setting (Sidik et al., 2016). The change of representation of guide DNA sequences in the population after three rounds of invasion and growth was translated into a fitness score based on a log_2_ scale. An analysis of the 68 glycogenes in Table 1 revealed a range of fitness values (Fig. 1) that roughly superimposed on the range of fitness values of all genes. 19 had fitness values <-2.5, suggesting that they may be important for viability. This group was enriched for predicted functions in GPI-anchor assembly, and deficient in genes associated with *O*-glycosylation in the secretory pathway and glycosylation in the nucleocytoplasmic compartment.

### Glycogene disruption strategy

To associate glycogenes with assembly of the glycome and cellular activities, we investigated consequences of glycogene disruption. As summarized in Fig. 2, we implemented a variation of the double CRISPR/Cas9 approach described by Shen et al. (2014), in which two gRNAs are employed with the intent of replacing most of the coding region with the pyrimethamine resistance marker in natively Ku80-positive parasite strains (Long et al., 2016). gRNA sequences were chosen using the Eukaryotic Pathogen CRISPR guide RNA/DNA Design Tool (Peng and Tarleton, 2015). We modified a previously constructed transient expression plasmid designed to express a single gRNA sequence and the Cas9 endonuclease to express 2 gRNAs as shown in Fig. 2E. Transient expression of the plasmid, together with a universal DHFR amplicon prepared by a simple PCR reaction, results in two double stranded breaks that can be repaired via NHEJ by the insertion of an amplicon encoding DHFR cassette. Treatment with pyrimethamine yields parasites that have incorporated the amplicon, many of which have insertions at the Cas9 cleavage sites. Disruption was confirmed by PCR experiments utilizing oligonucleotide primer pairs that flank both gRNA target sites, and primer pairs that bridge from outside the expected deletion and either end of the DHFR cassette, as outlined in Fig. 2F. This approach has the benefit of using a universal DHFR amplicon and does not stress the cells due to the absence of NHEJ (Ku80), which is commonly used for molecular biology studies of *Toxoplasma*. In addition, this method can in principle be applied to any wild-type clinical isolate regardless of the family type. We applied that double-CRISPR/Cas9 disruption method to selected genes representing each of the major glycan classes but whose fitness scores (Fig. 1) did not suggest inviability, as summarized in Table 1.

Of the 68 glycogene candidates, dual gRNA plasmids were constructed for 30, and single gRNA plasmids were generated for 4 others (summarized in Table S1). Of the 30 for which pDG constructs were available, 17 were used to modify the target gene, and 8 were unsuccessful, as determined by PCR studies (Table S1). Five genes were not tested, usually because mutants became available from other studies during the course of this project. Of the eight unsuccessful trials, each was independently repeated, with the same outcome, and three were re-tried with new pairs of gRNAs, with the same negative outcome (Table S1). This was unexpected based on fitness scores. Of the 17 successful disruptions, ten were full deletions engaging both gRNA sites, and seven resulted in effects at only one of the gRNA sites. Fifteen of these were examined glycomically to address their function and to assess whether the genetic disruption resulted in a loss of function (see below). They were also examined for cell level effects using a plaque assay in the next section.

Each transfection generated a spectrum of genetic lesions, which were evaluated by PCR trials involving primers that hybridized with gene-specific flanking sequences and DHFR specific sequences. Clones were classified based on evidence for a double cut, in which the intervening coding region was deleted and replaced with the DHFR cassette and/or other DNA, and single cut at either the 5’- or 3’-gRNA site and insertion of the DHFR and/or other DNA, or only an improper repair with no introduced DNA, as summarized for each gene in Table S1. Most if not all clones analyzed for individual genes showed evidence consistent with incorporation of the DHFR resistance cassette at one or both of the double stranded break sites. An expanded analysis of the results from three representative gene loci (*galE*, *spy* and *pofut2*) is presented in Figures S2-S4.

### Assessment of cellular consequences

To examine cellular effects of disrupting a glycogene, we employed a 6-d monolayer infection model, in which spreading infection from an initial single cell generates a lytic plaque, whose area is a measure of successful cycles of parasite invasion, proliferation and egress. As shown in Fig. 3, nine of the 13 strains examined exhibited significant deficiency in this setting. These included mutations predicted to affect *N*- or *O*-glycosylation in the secretory pathway, *O*-glycosylation in the nucleocytoplasm, or sugar-nucleotide precursor supply, as detailed below. The effects of nine of the disruptions reasonably correlated with fitness scores from Sidik at al. (Fig. 1), but diverged for four genes. The effects of three disruptions were more severe than observed in the fitness assay. Though this might be attributable to unknown spurious genetic changes, it is notable that these three genes are associated with protein fucosylation: synthesis of GDP-Fuc or its utilization by the Spy or POFUT2 glycosyltransferases (see below). Thus the variations might be due to the different parasite activities assayed as, in contrast to the plaque assay, the fitness assay did not test parasite egress from host cells. Though these correlations strongly implicate the importance of different glycosylation pathways for overall parasite success in cell culture, follow-up studies beyond the scope of this resource will be required to confirm causality for individual glycogenes.

**Fig. 3.**
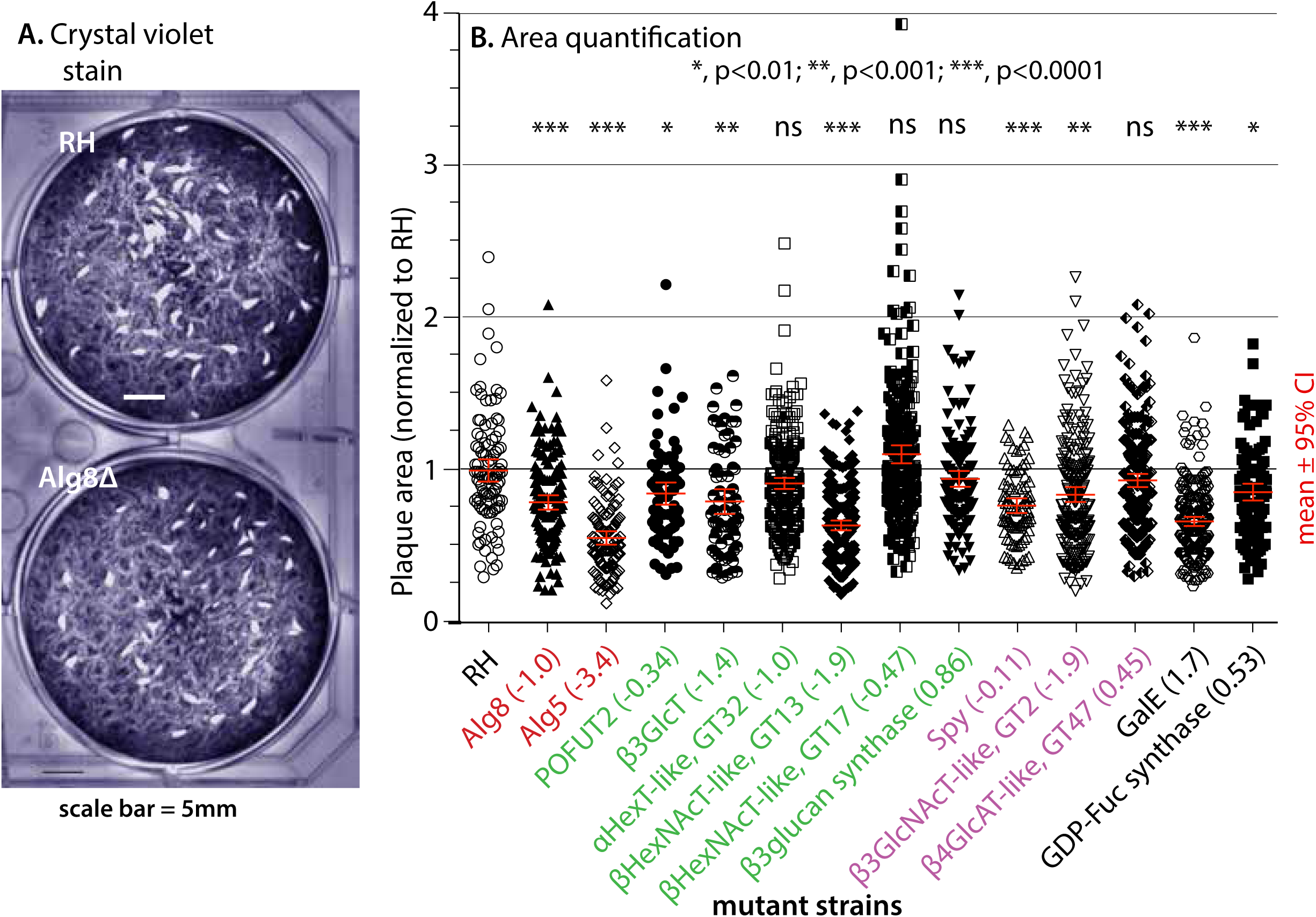
Phenotype analysis. A clone of each strain was analyzed using a plaque assay on confluent human foreskin fibroblast (HFF) monolayers, and plaque areas were measured 6 d after clonal infection. (**A)** Image comparing plaques from RH and *alg8*Δ strains. (**B)** Dot plots from pairwise comparisons of mutant and parental (RH) strains. Gene names are color-coded to indicate their glycosylation pathway association (red, *N*-glycans; green, other secretory pathway glycans; purple, cytoplasmic glycans; black, precursor assembly), and Fitness scores from Fig. 1 are indicated in parentheses. To facilitate comparisons between trials, the mean areas of RH plaques were normalized to one and the respective mutant plaque areas scaled accordingly (only one representative RH experiment is graphed for clarity). Mean values ± 2 standard deviations (95% confidence interval) are shown with red bars. Statistical significance is shown at the top. Data were pooled from 3 independent trials, which each independently conformed to the same trend.

### Assessment of glycomic consequences

In order to assess the biochemical consequences of glycogene disruption, we adapted established methods to evaluate the parasite glycome. Since the parasite grows within human host cells, which are themselves reared in the presence of bovine serum, measures are required to minimize contamination with human and bovine glycans. Thus we allowed tachyzoites to spontaneously lyse out of fibroblast monolayers in low (1%) serum medium, and washed the parasites by centrifugation with serum-free medium. Since glycans are found in glycolipids and glycoproteins, and as polysaccharides, which vary in their chemical characteristics, we adapted a classical scheme of Folch extraction to generate fractions enriched in glycolipids, GPI-anchored glycoproteins, and other glycoproteins, and then applied chemical or enzymatic methods to release glycans from these fractions to be analyzed by mass spectrometry (Fig. S5). The various chemical linkages of glycans to protein amino acids necessitate specialized methods for their release. In general, *N*-glycans are released enzymatically using PNGaseF, *O*-glycans are released chemically using NaOH to induce β-elimination, and GPI-anchor glycans are released using aqueous HF that cleaves phosphodiester linkages. For monosaccharide glycans that are too short to recover by current methods, we Western-blotted the carrier glycoproteins with antibodies (Abs) and lectins; alternatively, glycans might be analyzed as glycopeptides by MS after trypsinization.

### *N*-glycans

*N*-glycans are present on dozens if not hundreds of proteins that traverse the *Toxoplasma* secretory pathway (Fauquenoy et al., 2011; Fauquenoy et al., 2008; Luo et al., 2011), though the density of *N*-sequons to which they are linked is less than in most eukaryotes (Bushkin et al., 2010). We identified genes expected to assemble a Glc_3_Man_5_GlcNAc_2_-P-moiety on the P-Dol precursor (Table 1), in accord with previous studies (Samuelson et al., 2005). *N*-glycans are typically extensively processed after transfer to the protein, but only enzymes that remove the Glc residues were detected in *Toxoplasma* (Table 1). Notably, the processing α-mannosidases and UDP-Glc-dependent α-glucosyltransferase, found in most eukaryotes, are absent. Together with the absence of other genes likely due to secondary loss, *N*-glycans are not expected to have a role in chaperone assisted protein folding and quality control (Banerjee et al., 2007; Samuelson and Robbins, 2015). Studies indicate that *N*-glycosylation is important for parasite motility and invasion (Fauquenoy et al., 2011; Luk et al., 2008), consistent with the low fitness scores (<-2.5) for 4 of the 5 enzymes that assemble the Man_5_GlcNAc_2_-core (Fig. 4A).

**Fig. 4.**
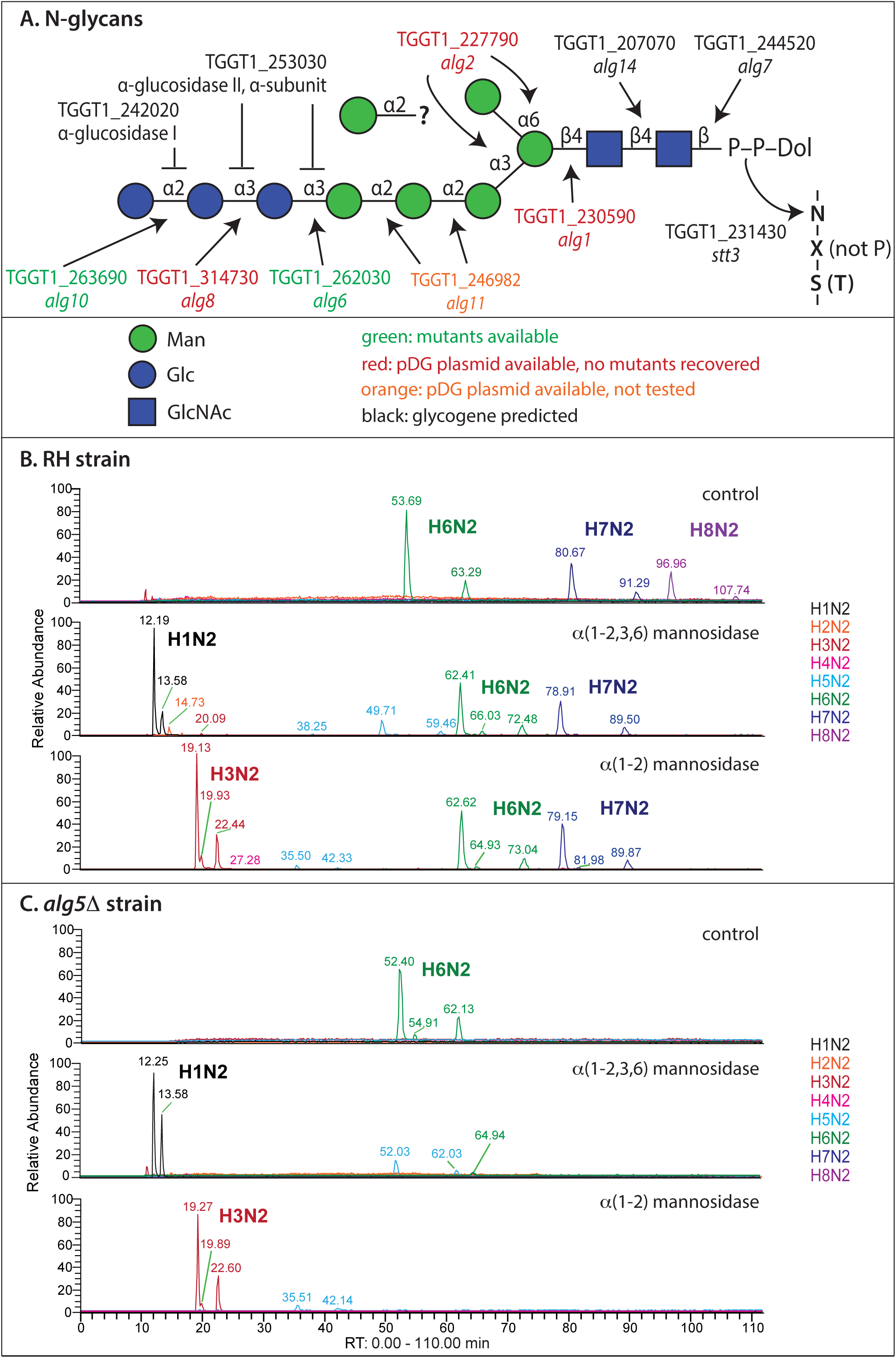
*N*-glycan glycogenes and nanoLC-MS analysis of parasite *N*-glycans. (**A**) The *N*-glycan precursor is assembled on dolichol-pyrophosphate, one sugar at a time from right to left. Genes that contribute to biosynthetic steps are depicted. The status of gene editing is color-coded according to the key. The complete Glc_3_Man_6_GlcNAc_2_ glycan is transferred *en bloc* to protein Asn side chains in an NxS/T motif by *stt3* and variably trimmed by α-glucosidases-I and –II (Table 1A). Glycans are represented according to the Consortium for Functional Glycomics system of nomenclature (Varki et al., 2015). (**B, C**) *N*-glycans were prepared by treating tryptic digests of total parasite protein powders with PNGaseF. The released glycans were permethylated and subjected to nLC on a C18 column followed by MS^n^ in a linear ion-trap MS. Extracted ion chromatograms (EIC) for all detected *N*-glycan-like, lithiated compositions (H1N2 through H8N2) species are shown before and after digestion with jack bean α-mannosidase (α1-2/3/6) or *Aspergillus satoi* (α1-2) mannosidase to probe the identity of the hexose (H) species. The two peaks observed for each species are α- and β-anomers. Minor related *N*-glycan-like species are shown in Fig. S6A. (**B**) RH (parental). (**C**) *alg5*Δ-mutant, which is unable to form Dol-P-Glc, the Glc donor for terminal α-glucosylation.

nLC-MS(n)-based glycomic profiling was employed to interrogate consequences of *N*-glycogene disruptions. Analysis of unfractionated tachyzoites identified three prominent *N*-glycan-like species: H8N2, H7N2 and H6N2 (Fig. 4B), and other minor species including H9N2, H5N2 and H4N2 (Fig. S6A), where H indicates the number of hexose and N the number of HexNAc residues. H8N2 was initially expected to represent Glc_3_Man_5_GlcNAc_2_, with H7N2 and H6N2 being processed forms lacking one or two Glc residues. The absence of complex, hybrid and paucimannose *N*-glycans confirmed minimal contamination from host or serum glycans (Fig. S6B). However, the H6N2 species elutes at the same time both in RH and in a mutant disrupted in *alg5* (Fig. 4), the enzyme that generates the Dol-P-Glc precursor required for glucosylation. H7N2 and H8N2 were absent from *alg5*Δ cells, suggesting that they represented H6N2 containing one or two Glc caps, respectively. To address the identity of the Hex residues, the samples were treated with jack bean α-mannosidase, which cleaves α-1-2/3/6-linked mannoses. H6N2 from both RH (parental) and *alg5*Δ cells, which are likely identical owing to co-elution of both of their anomers from the C18 column, were both converted to H1N2 (Fig. 4B, C), indicating that the extra Hex residue is an αMan. Furthermore, α2-mannosidase converted the H6N2 to H3N2, indicating that the αMan is 2-linked. After treatment with either α-mannosidase, H8N2 appeared to convert to H7N2 (similar elution time as the original H7N2), and H7N2 to convert to H6N2 (different elution time compared to original H6N2), which suggests that the extra α2Man occurs in both of these species. This conclusion is based on the assumption that jack bean α-mannosidase, at the enzyme concentration used, can only access the α6Man residue after the α3-arm is partially trimmed, as can only occur in the absence of the αGlc caps.

In this new model, we propose that *Toxoplasma* assembles an unusual Glc_3_Man_6_GlcNAc_2_-linked precursor (Fig. 4A), and that the 6^th^ Man residue is α2-linked to the α6Man, though linkage to the α3Man or to the non-reducing end α2Man cannot be excluded. The responsible αManT is unknown, but might be Alg11, which applies the other two α2Man before the precursor flips from the cytoplasm to the lumen of the rER, or might be attached after flipping by the GPI-anchored pigB, which also applies an α2Man to an α6Man. This model is consistent with Hex_9_GlcNAc_2_ being the largest and most abundant lipid-linked oligosaccharide in *Toxoplasma* (Garénaux et al., 2008). After transfer to protein by the oligosaccharyltransferase (Shams-Eldin et al., 2005), partial trimming down to H6N2 is consistent with evidence for α-glucosidase-I and α-glucosidase-II genes (Fig. 4A, Table 1A). The minor H9N2, which was inconsistently observed, is evidently converted to H8N2 by jack bean α-mannosidase, and the minor H5N2 and H4N2 species were eliminated by treatment with jack bean α-mannosidase, indicating that they are canonical Man_5_GlcNAc_2_ and Man_4_GlcNAc_2_ species that might have resulted from processing by lysosomal hydrolases, as described in *Dictyostelium* (Hykollari et al., 2014). We detected no effect of *alg10*Δ, expected to assemble the 3^rd^, α2-linked, Glc, consistent with its efficient removal by α-glucosidase-1 (Fig. 4A), as expected. Our analysis of the putative *alg8*-mutant yielded a reduced level of H8N2 relative to H7N2 and H6N2 and a small plaque phenotype (Fig. 3). This difference suggested the genetic lesion, which consisted of an insertion of DHFR in a poorly conserved region of the 5’-end of the protein coding region, is likely a hypomorphic rather than a null allele. We found no evidence for host derived *N*-glycans (Garénaux et al., 2008), but this could be the result of our use of spontaneously lysed extracellular parasites, which may have turned over host-derived glycans.

Thus, *Toxoplasma* proteins appear to be decorated with just three major species of *N*-glycans, each with an unexpected and unusual structure: Glc_2_Man_6_GlcNAc_2_, Glc_1_Man_6_GlcNAc_2_, and Man_6_GlcNAc_2_ (Fig. 4A). *N*-glycans of *Cryptosporidium*, a related apicomplexan, include just two forms, Hex_6_HexNAc_2_ and Hex_5_HexNAc_2_, each of which lacks an extension on the short mannose arm and so likely represents Glc_1_Man_5_GlcNAc_2_ and Man_5_GlcNAc_2_, respectively (Haserick et al., 2017). Failure to trim the Glc residues is consistent with the absence of the quality control mechanism, in which they participate in other eukaryotes (Bushkin et al., 2010; Samuelson and Robbins, 2015). Attempts to genetically disrupt assembly of the Man_6_GlcNAc_2_-portion of the *N*-glycan precursor were unsuccessful, consistent with essentiality of its highly conserved Man_5_GlcNAc_2_ core. Disruption of assembly of the terminal αGlc residues, which are retained on many *N*-glycans, was deleterious for the parasite. Thus the αGlc termini contribute functionality to *Toxoplasma N*-glycans, which can in the future be investigated using the respective pDG plasmids described here.

### *O*-glycan glycogenes associated with the secretory pathway

*O*-glycans are also found on dozens if not hundreds of glycoproteins that are processed via the parasite rER and Golgi based mainly on lectin capture studies (Luo et al., 2011; Wang et al., 2016). The genome encodes five pp-αGalNAcTs (Table 1C and Fig. 5A) that initiate mucin-type *O*-glycan assembly (Stwora-Wojczyk et al., 2004), i.e., glycans that are linked via αGalNAc-residues to Thr/Ser residues that are often clustered in hydrophilic sequence repeats. In addition, two genes (Table 1C, Fig. 5B) are predicted to assemble a second type of *O*-glycan (Glcβ1,3Fuc1α-) involved in quality control of thrombospondin type 1 repeat-domains (TSR) and protein-protein interactions in animals (Vasudevan and Haltiwanger, 2014). Four Golgi-associated GTs of unknown function are also predicted (Table 1E and Fig 5D). With fitness scores ranging from -1.9 to 1.4 (Table 1), the contributions of these genes to tachyzoite biology is predicted to be modulatory rather than essential for viability. Mutational analyses of the pp-αGalNAcTs and a sugar nucleotide transporter that may selectively impact *O*-glycan assembly, indicate functional roles in the integrity of semi-dormant bradyzoite cyst walls (Tomita et al., 2017), which contains mucin type glycoproteins including Cst1 (Tomita et al., 2013), and tissue cyst persistence in mice (Caffaro et al., 2013).

**Fig. 5.**
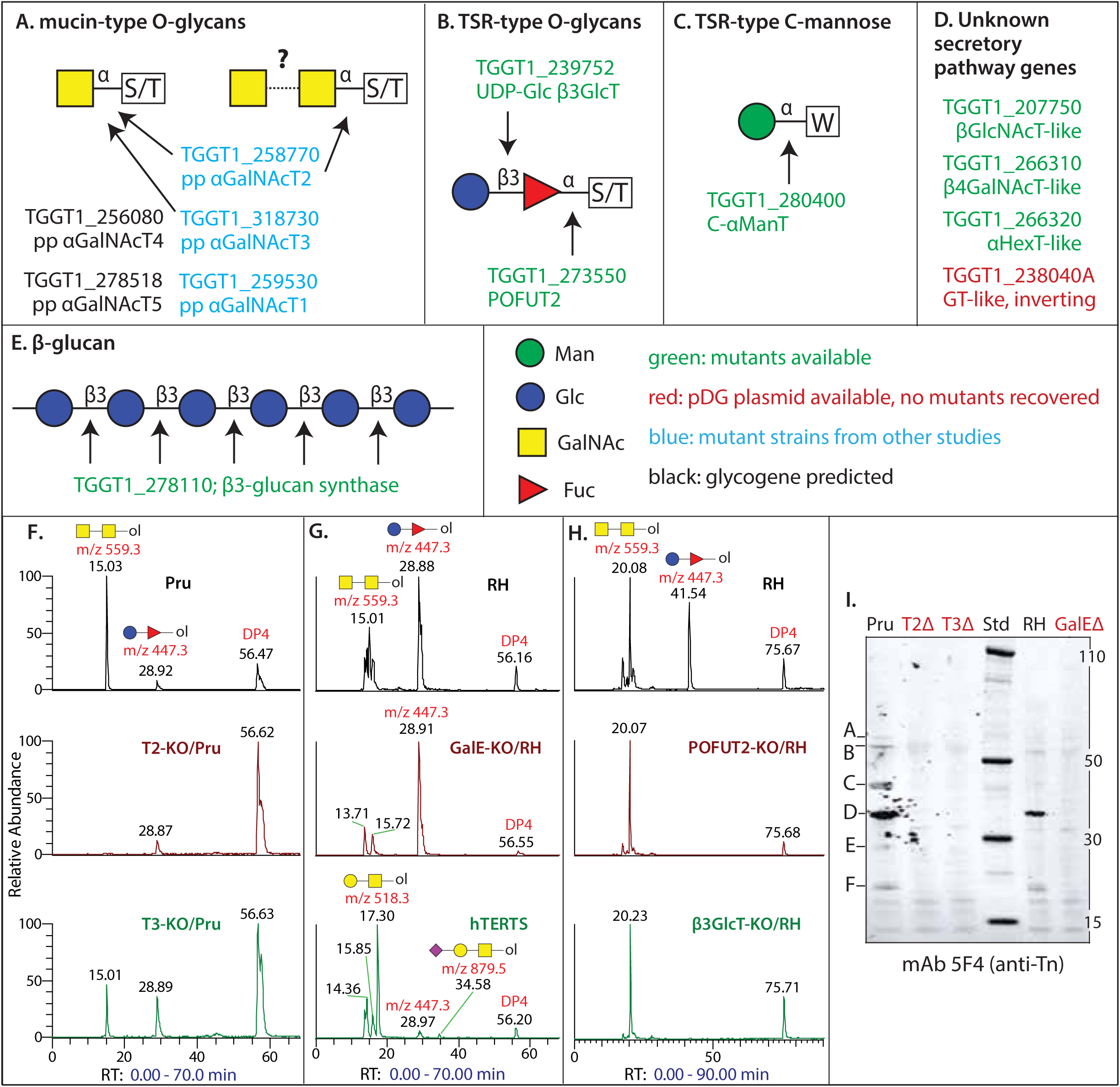
*O*-glycan, *C*-mannose and polysaccharide glycogenes in the secretory pathway and their glycomic analysis. (**A**) Mucin-type *O*-glycans. Tn antigen (GalNAcα-Ser/Thr) and the N2 disaccharide, consisting of a HexNAc of uncertain identity (potentially GalNAc) linked to αGalNAc, inferred from MS glycomic and lectin labeling studies. Five pp GalNAcTs are predicted to catalyze formation of the Tn antigen, and T2 and T3 are confirmed. T2 is required for accumulation of N2, but the enzyme responsible for the second sugar is unknown, except that genes in green in panel D are not required. (**B**) TSR-type Glc-Fuc– disaccharide and its documented GT genes (see Fig. 7C). (**C**) GT for *C*-mannosylation. (**D**) Four genes encoding predicted sugar nucleotide dependent GT that appear to be type 2 membrane proteins in the secretory pathway. The glycans to which they contribute are yet to be identified. (**E**) GT for assembly of the oocyst wall β3-glucan, whose priming mechanism is unknown. (**F-H**) *O*-glycans recovered reductive β-elimination, permethylated, and subjected to nLC on a C18 column followed by MS^n^ in a linear ion-trap MS as in Fig. 4. EICs for N2 (drawn as GalNAc-GalNAc-) and HdH (drawn as Glc-Fuc-) species (lithiated adducts ± 70 ppm), and the DP4 standard, are shown. (**F**) Parental strains (Type 2 Pru) and two derivative mutant strains (pp-αGalNAcT2 and pp-αGalNAcT3). (**G**) The parental strain (Type 1 RH) and the derivative *galE*Δ strain, and an extract of host hTERT cells, showing HN, HNSa and HdH, but lacking N2. (**H**) RH and two derivative mutants predicted to encode the sequential addition of αFuc (*POFUT2*Δ) and βGlc (*β3GlcT*Δ). (**I**) Western blot analysis of total RH strain tachyzoites for the Tn-antigen, probed with mAb 5F4. Similar results were obtained with mAb 1E3, except that mAb-reactive proteins were differentially emphasized.

Little is known about *Toxoplasma O*-glycans, so we initiated a global *O*-glycomic analysis of material released from total tachyzoite protein powder by classical reductive β-elimination (Fig. S7). Remarkably, only two *O*-glycan types could be confidently assigned, with compositions of N2 (HexNAc2) and HdH (Hex-deoxyHex). The assignments were verified by exact mass and diagnostic MS^2^ and MS^3^ fragment ions. These ions did not appear to be host derived because i) N2 was not detected in host cell samples, ii) HdH is more abundant relative to other glycans in *Toxoplasma* vs. host cell samples, and iii) host O-glycans, including HN, H2N, H3N, and sialylated derivatives (Fig. 5G and not shown), were not detected in the *Toxoplasma* sample (Fig. S7A). The N2 and HdH ions were detected at higher levels than the Man_6_GlcNAc_2_ *N*-glycan that is inefficiently released by conditions of β-elimination. Monosaccharide modifications cannot be confidently assigned by this method. These findings indicate a limited repertoire of *O*-glycosylation at the proliferative tachyzoite stage and document the selectivity of the method for parasite glycans.

The N2 species is potentially representative of a mucin-type *O*-glycan and, based on studies using the lectins jacalin, *Vicia villosa* agglutinin, *Dolichos biflorus* agglutinin, and *Helix pomatia* agglutinin, expression of a GalNAc-GalNAc-structure was previously proposed (Fig. 5A). Analysis of the *O*-glycomes of pp-αGalNAcT2 (*gant2*Δ) and pp-αGalNAcT3 (*gant3*Δ) mutants showed that N2 depended on Gant2, with no effect observed on the expression of the HdH species (Fig. 5F). N2 was detected in *gant3*Δ extracts, though its apparent abundance relative to HdH ion was consistently reduced. Furthermore, N2 was also dependent on the expression of *galE* (see below), the epimerase that generates the sugar nucleotide UDP-GalNAc precursor for the pp-αGalNAcTs from UDP-GlcNAc (Fig. 5G). These results indicate that the reducing end N residue is GalNAc, but do not reveal the identity of the second N. The involvement of two genes predicted to encode Golgi HexNAcTs, TGGT1_266310 and TGGT1_207750 (Table 1E), have been excluded based on analysis of their *O*-glycans (Fig. S7 and data not shown). Thus tachyzoites express only one major mucin-type *O*-linked oligosaccharide, a disaccharide that may correspond to the previously predicted core 5 structure (GalNAcα1,3GalNAcα1-Thr). This prediction was based on lectin studies on bradyzoites (Tomita et al., 2017), which matched findings from metabolic labeling of a disaccharide on the GRA2 glycoprotein (Zinecker et al., 1998) that can be interpreted as GalNAc-GalNAc-based on more recently available information that the jacalin lectin can recognize non-reducing terminal GalNAc as well as Gal (Tachibana et al., 2006).

To investigate the potential expression of non-extended αGalNAc, referred to as Tn-antigen, we utilized anti-Tn mAbs developed from studies on human cancer (Steentoft et al., 2011). As shown in Fig. 5I, using two validated (see Methods) anti-Tn mAbs, we observed that both type 1 (RH) and type 2 (Pru) strains expressed at least 6 proteins (A-F) carrying the Tn-antigen in the *M*_r_ range of 20,000-60,000. Proteins in this *M*r range have also been detected with various GalNAc-preferring lectins (Tomita et al., 2017; Wang et al., 2016). mAb recognition was dependent on both Gant2 and Gant3, as well as, as expected, GalE. Since only one pp-αGalNAcT is required to express Tn-antigen, these findings support the model of Tomita et al. (2017), consistent with precedent in animals, that Gant2 is a priming pp-αGalNAcT, and Gant3 is a follow-on pp-αGalNAcT that adds to the number of structures. Since N2 is more dependent on Gant2 than on GAnt3, N2 is likely to be assembled on the amplified number of Tn-antigen structures generated by Gant3.

The HdH ion is a candidate for the Glcβ1,3Fucα1-disaccharide that has been reported to be linked to Ser/Thr residues at consensus positions of TSR domains in animal glycoproteins (Vasudevan et al., 2015), and of *Plasmodium* CSP and TRAP proteins that also have the consensus peptide motif (Swearingen et al., 2016). MS^2^ and MS^3^ fragmentation indicated that the order of the sugars is Hex-dHex-(data not shown). Disruption of the *Plasmodium* homolog of *pofut2*, which catalyzes addition of αFuc to Ser/Thr, prevents assembly of the glycan (Lopaticki et al., 2017). Similarly, our *pofut2*Δ strain also blocked accumulation of HdH (Fig. 5H), demonstrating by analogy that the core deoxyHex residue is αFuc. Two recent preliminary reports confirm this result in *Toxoplasma* (Bandini et al., 2018; Khurana et al., 2018). The genome harbors a CAZy GT31 gene (TGGT1_239752 or *glt2*; Table 1) with homology to animal B3GlcT, which catalyzes addition of the βGlc cap at the 3-position of Fuc. Our *glt2*Δ strain also failed to express the disaccharide (Fig. 5H), confirming its role in its assembly, and indicating conservation of the Glcβ1,3Fucα1-disaccharide in *Toxoplasma* (Fig. 5B). Both assembly mutants exhibit a small plaque phenotype (Fig. 3), suggesting conservation of a role in quality control of folding of the several GRA proteins that bear the consensus motif in their TSR-repeats (Vasudevan et al., 2015).

In this first investigation of its *O*-glycome by mass spectrometry, *Toxoplasma* tachyzoites are found to exhibit a limited repertoire of *O*-linked oligosaccharides, just two disaccharide species that can be detected at the whole cell level. The precursor for the GalNAc-GalNAc-species, Tn-antigen (GalNAc-), also accumulates at detectable levels, and *O*-Fuc was also recently detected on TSR5 of MIC2 in a glycopeptide analysis (Bandini et al. 2018). Phylogenetic analyses of the genes known to assemble *Toxoplasma O*-glycans, including the Tn antigen, Glcβ1,3Fucα1-, and *C*-Man described below, are found in many other apicomplexans but more related to their metazoan counterparts than genes found in other protists (unpublished). This is good evidence for the horizontal transfer of *O*- and *C*-glycosylation mechanisms from the genome of a host cell, within which apicomplexans are obligate residents.

### GPI-anchor and GIPL-glycogenes

Numerous parasite proteins are linked to the cell surface via glycophosphatidylinositol (GPI) anchors. The glycan portion of *Toxoplasma* GPI anchors conforms to the conventional Man_3_GlcNH_2_inositol core formed by most eukaryotes but are reportedly modified at the core Man by a 3-linked βGalNAc residue (Azzouz et al., 2006). Some GPI anchor precursors also remain free as GIPLs, and are distinguished by an additional αGlc 4-linked to the βGalNAc (Fig. 6A). In addition, there is strain variation with regard to the presence of the phosphoethanolamine and αGlc termini in GIPLs (Niehus et al., 2014). Besides serving a structural role, GPI anchors and possibly GIPLs are recognized by host cell TLR2 and probably galectins as co-receptors (Debierre-Grockiego et al., 2010). Studies targeting either the GPI anchor or the galectin implicate their participation in the infection of host cells. The novel Glcα1,4GalNAcβ1-branch has been the subject of vaccine candidate and antibody-based diagnostics (Anish et al., 2014).

**Fig. 6.**
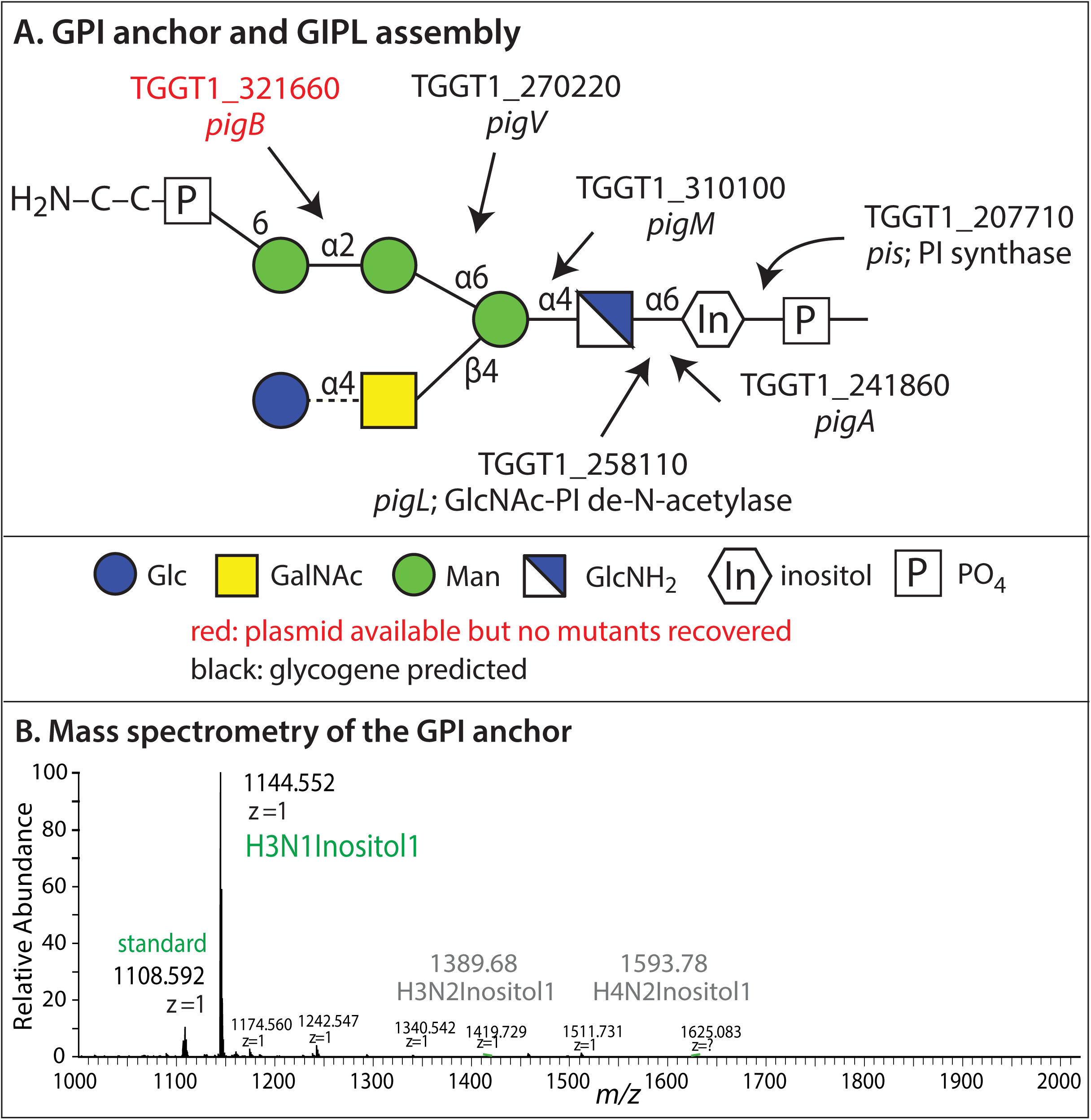
GPIL and GPI-anchor biosynthesis glycogenes. (**A**) Free GIPLs and GPI-anchors share a related glycan linked to a diacylglycerol on the right. Glycan assembly proceeds stepwise from right to left along the upper arm, followed by the lower arm (Azzouz et al., 2006), by the enzymes predicted. The GPI-anchor precursor receives only the lower arm β4GalNAc, whereas the GPIL also receives an α4Glc (dashed line). Phosphoethanolamine is then conjugated to the terminal α2Man, and the resulting NH_2_-group of the GPI precursor is conjugated to the polypeptide C-terminus by a multisubunit transamidase. Potential fatty acyl modifications are not shown. (**B**) The glycan region of GPI-anchored proteins enriched from RH strain tachyzoites was released after saponification by HF, which cleaves phosphodiester bonds, subjected to re-N-acetylation and permethylation, and analyzed by MS^n^. *m/z* values corresponding to glycans containing one or two of the sugars of the predicted lower arm are indicated in gray, but were not detected in this study.

We developed a new MS-based method to analyze GPI anchor glycans released from an enriched pool of GPI-anchored proteins, by saponification and HF to break the phosphodiester linkages at both ends of the glycan. Unexpectedly, our analysis of strain RH tachyzoites yielded only the linear Man_3_GnIn core lacking a side chain (Fig. 6B).

The high degree of conservation of the enzymes that assemble the canonical Man_3_GnIn backbone enabled confident prediction of the enzyme coding genes (Table 1B). The phosphatidylinositol synthase (*pis*) that assembles the initial precursor from CDP-diacylglycerol and myoinositol has been described (Seron et al., 2000). *pis*, the *pigA* GlcNAcT, and the *pigM*, *pigV*, and *pigB* ManTs have very low fitness values, -3.1 to -5.1 (Fig. 1), indicating essentiality of the complete backbone. Our attempts to disrupt the final αManT (*pigB*) of the pathway were unsuccessful (Fig. 6A), consistent with the fitness data. With respect to the GTs that mediate the formation of the disaccharide side chain, the *Toxoplasma* genome encodes two βHexNAcT-like GTs that are predicted to be Golgi-associated type 2 membrane proteins, TGGT1_266310, TGGT1_207750, and one predicted αHexT that is a type 2 membrane protein, TGGT1_266320. These candidates, which have modest fitness values, were each successfully disrupted in our RH strain, but their effects on the assembly of the GPI-anchor glycan branch could not be evaluated owing to its absence in our studies (Fig. 6B).

In this first examination of *Toxoplasma* GPI-anchor glycans by mass spectrometry, we confirmed the core backbone, but did not detect a previously described unique glycan feature. Nevertheless, we have enumerated a plausible genetic basis for the assembly of the glycan part of the anchor and GIPLs with their branch chains, and developed tools for their analysis in future studies in other strains and under other conditions.

### Other glycogenes associated with the secretory pathway

In addition to *N*- and *O*-glycosylation of secretory proteins, and their C-terminal glycolipidation, *Toxoplasma* also modifies a C-atom of the side chain of certain Trp residues with αMan (Fig. 5C). Originally found in association with a multi-Trp sequence motif in animal TSRs, this monosaccharide modification and the motif are conserved on CSP and TRAP proteins in *Plasmodium*, a related apicomplexan (Vasudevan et al., 2015). Both *Plasmodium* and *Toxoplasma* genomes encode a high scoring homolog of Dpy19 (Table 1D) that mediates formation of this linkage from Dol-P-Man in worms, and the *Toxoplasma* enzyme was recently shown to harbor the same activity (Hoppe et al., 2018). Using our double CRISPR plasmid we achieved a single insertion near the C-terminus of the predicted protein (Fig. 5C), but it is unclear whether this affects its C-αManT activity. Its -2.4 fitness score is consistent with importance for growth, potentially related to quality control of secretion of the proteins carrying TSRs, possibly in concert with Glc-Fuc-modifications (Scherbackova et al., 2017). Since there is no method known to release C-Man, monitoring the outcome will depend on glycopeptide analyses beyond the scope of this project.

The oocyst wall, formed during the sexual cycle in the gut of the cat host, contains disulfide rich proteins and a β3glucan polysaccharide (Fig. 5E), and the genome encodes a predicted β3glucan synthase (Bushkin et al., 2012; Samuelson et al., 2013). Its disruption had no effect on proliferation of tachyzoites (Fig. 3), as expected because the polymer is assembled at another stage of the life cycle. Deletion of β3glucan synthase also has no effect on bradyzoite and tissue cyst formation *in vitro* (Bushkin et al., 2012). A previously described hint for the presence of chitin in the bradyzoite cyst wall is not supported by evidence for a chitin synthase-like sequence in the genome.

In addition to the free GIPLs, there is evidence for a class of glycosphingolipid-like molecules that can be metabolically labeled with tritiated Ser and Gal (Azzouz et al., 2002). They are so classified based on the resistance of the radiolabeled material to saponification and their sensitivity to known inhibitors of sphingolipid synthesis. The tritiated Gal was evidently incorporated as GalNAc, which might be mediated by one of the two predicted βHexNAcTs found in the genome (Table 1E).

Our GT predictions yield four unidentified genes (Table 1E, Fig. 5D), and our inventory of known glycans yields four linkages in need of GTs (N2, GPI-anchor, and glycosphingolipid). Future studies are needed to make the potential connections, and this summary does not exclude the possibility of lower abundance glycans or novel GT sequences yet to be discovered.

### Cytoplasmic glycosylation glycogenes

Proteins of the nucleocytoplasmic compartment of eukaryotic cells are also potentially subject to glycosylation (West and Hart, 2017). Because these involve different GT genes and often require detection by different methods, this class of glycosylation events is often overlooked.

A subunit of the SCF class of E3 ubiquitin ligases, Skp1, has been shown to be modified by an *O*-linked pentasaccharide (Fig. 7A). A homolog of this glycan was originally discovered in the social amoeba *Dictyostelium* by metabolic labeling with [^3^H]fucose, and subsequently defined by mass spectrometry and NMR (Sheikh et al., 2017). The glycan is linked in alkali-resistant fashion to a hydroxyproline, which therefore must be analyzed as tryptic glycopeptides, and may be restricted to only a single protein and so is not abundant. The *Toxoplasma* genome contains homologs of two of the GT genes required to assemble the *Dictyostelium* glycan (Rahman et al., 2016), which led to the discovery of a related pentasaccharide in *Toxoplasma* that is assembled by the two related and two unrelated GTs (Rahman et al., 2017). The Skp1 GT genes are not essential, but their disruption by conventional double crossover homologous recombination each generated small plaques, consistent with a role for the glycan in O_2_-sensing originating from the O_2_-dependence of the prolyl hydroxylase that forms the Hyp residue on which it is assembled (Xu et al., 2012; West and Blader, 2015). pDG plasmids and mutants for the first and final GTs in the pathway, *gnt1* and *gat1* (Table 1G), are available.

**Fig. 7.**
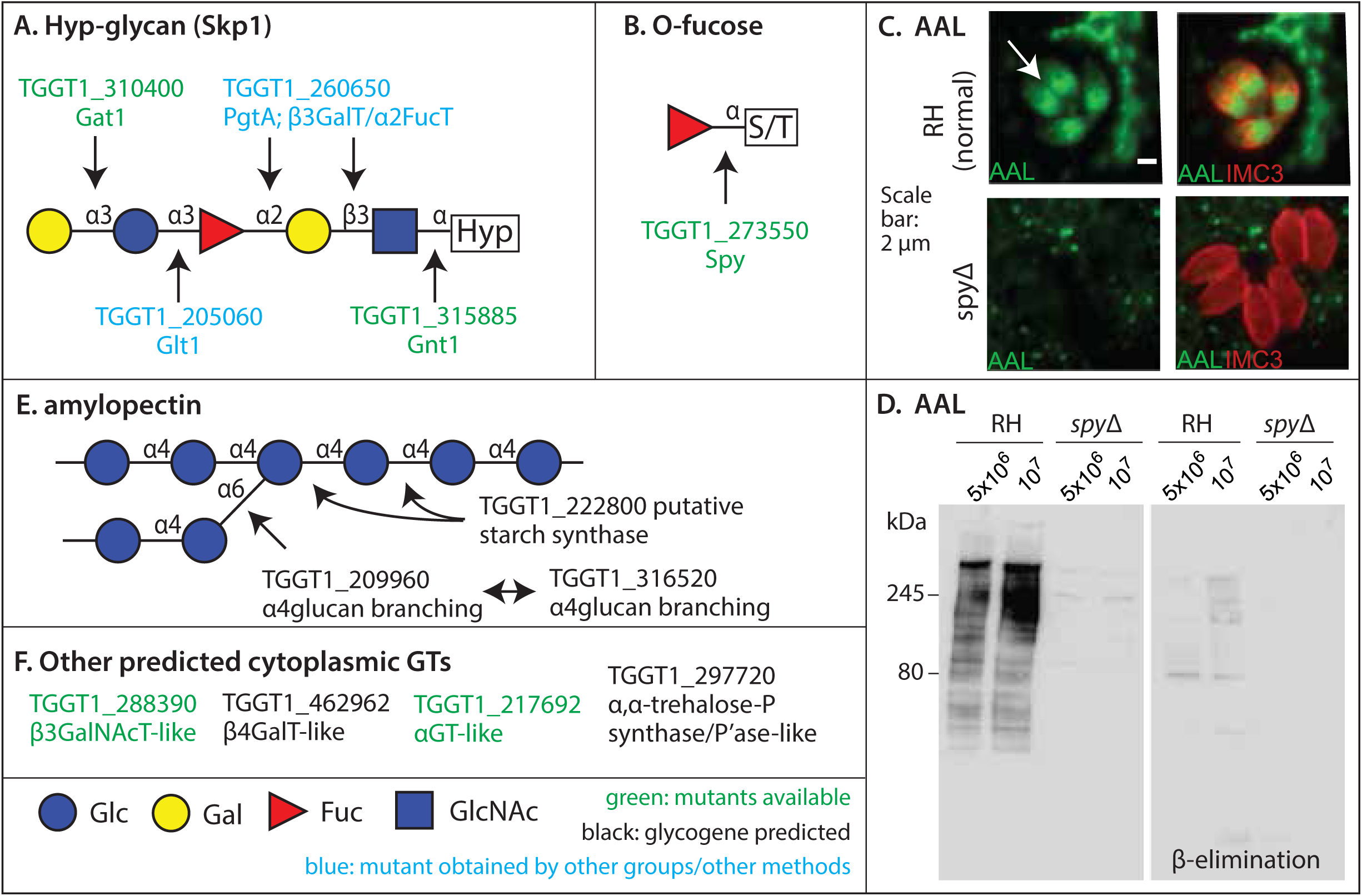
Cytoplasmic GT glycogenes. (**A**) Assembly of the hydroxyproline-linked pentasaccharide on the E3^SCF^Ub-ligase subunit Skp1. (**B**) Formation of *O*-Fuc on Ser/Thr residues of nucleocytoplasmic proteins. AAL (green) binding to 4 nuclei of parental RH strain parasites (white arrow, upper left), housed within the same parasitophorous vacuole within a host fibroblast, is lost in the six *spy*Δ cells shown (lower left). To the right, IMC3 (red) labeling localizes to the parasite inner membrane complex whose organization varies according to the cell cycle. (**D**) AAL labeling of Western blots of parental strain whole cell extracts is lost in *spy*Δ cells. Subjecting the blotted membrane to conditions of β-elimination removed the majority of labeling, consistent with its *O*-linkage. (**E**) Genes associated with amylopectin assembly. Genes associated with its disassembly are in Table 1I. (**F**) Other genes predicted to encode sugar nucleotide-dependent cytoplasmic (or nuclear) GTs.

Recently, *Toxoplasma* nuclei were shown to be enriched in a novel Fuc modification that could be detected cytologically with the lectin *Aleuria aurantia* (AAL), which glycoproteomic analysis showed consisted of αFuc-modifications of Ser/Thr residues (Fig. 7B) often in clusters (Bandini et al., 2016). Our analysis predicted four unknown cytoplasmic (or nuclear) GTs, one of which was homologous to the *O*-βGlcNAcT (OGT or Secret agent) that modifies Ser/Thr residues of metazoan and higher plant nucleocytoplasmic proteins with *O*-βGlcNAc. Disruption of this gene, more closely related to its paralog *spindly* (*spy*) in *Arabidopsis*, abolishes AAL labeling of nuclei (Fig. 7C) and of extracts analyzed by Western blotting (Fig. 7D), indicating that Spy is an αFucT (or OFT) of nucleocytoplasmic proteins. This finding parallels that of a recent report showing *Arabidopsis* Spy is a *bona fide* OFT (Zentella et al., 2017). Despite the similarity of the Spy and OGT (Sec) sequences, a phylogenetic analysis of their evolution indicates that they are related by an ancient gene duplication possibly in prokaryotes, and diverged their donor sugar substrates at least by the time they appeared in eukaryotes (West et al., 2004). Spy appears to contribute functionality to the parasite according to the small plaque size phenotype (Fig. 3), though it did not contribute to fitness in the high throughput assay.

The cytoplasm is also the site of accumulation of amylopectin (Fig. 7E), a polymer of 4-linked αGlc with periodically spaced 6-linked α4glucan chains (Guérardel et al., 2005). Amylopectin assembles into semicrystalline floridean starch granules that are highly induced during bradyzoite differentiation. The *Toxoplasma* genome contains sequences for UDP-Glc dependent amylopectin assembly and disassembly that relate more to fungal and animal glycogen metabolism than to the ADP-Glc dependent pathways in plants (Coppin et al., 2005). The branched polymer is assembled by a starch synthase and potentially two branching enzymes (Fig. 7E), and disassembled by a debranching enzyme, starch phosphorylase (Sugi et al., 2017) (Table 1I), and exoglycosidases (not listed). A gene with sequence similarity to glycogenin, an α4GlcT that primes the synthesis of the related polymer glycogen in yeast and animals, is the Skp1 glycosyltransferase Gat1, which might have been the evolutionary precursor of glycogenin (Mandalasi et al., 2018). The starch assembly genes do not evidently contribute much to tachyzoite fitness, consistent with the induction of starch accumulation in bradyzoites and oocysts. Thus this pathway was not targeted in this study.

The genome encodes a gene with an α,α-trehalose-6-P synthase (TPS)-like domain, and a domain with resemblance to trehalose-6-phosphatase (TPP), glycohydrolase GH77 (α4-glucanotransferase), and CBM20 starch binding domains (Table 1I). Genes for trehalose formation and turnover are widely distributed in eukaryotes (Avonce et al., 2006), but the absence of a candidate for trehalase, or evidence for trehalose in *Toxoplasma*, suggests a potential role for this gene in starch metabolism. The function of this gene, which evidently contributes to fitness (-2. 9), remains unassigned.

There are two reports of a glycolipid, characterized by a dihexosyl-moiety, possibly including a di-Gal epitope, on ceramide, diacylglycerol and acylalkylglycerol lipids (Botté et al., 2008; Maréchal et al., 2002). This is reminiscent of galactolipid type structures associated with plastids in plants, so they might potentially be associated with the apicoplast, the relict plastid that persists in *Toxoplasma*. The enzymatic basis for the addition of these sugars remains to be determined.

Three GT-like sequences predicted to reside in the cytoplasmic compartment (Table 1H, Fig. 7F) remain to be assigned. pDG plasmids (as well as mutants for TGGT1_288390 and TGGT1_217692) are available for their investigation, and disruption of the β3GlcNAcT-like exhibits a small plaque phenotype (Fig. 3). The detection of their presumptive glycan products might require specialized approaches to be detected, if they follow the example of the Skp1 glycan.

### Sugar precursor glycogenes

Almost all glycosyltransferases rely on a sugar nucleotide or a Dol-P-sugar as the high energy sugar phosphate donor for the transfer of the sugar to the acceptor molecule. The formation of these donors from metabolic precursors tends to be well conserved. Fig. 8A-C summarize the metabolic pathways proposed to generate these compounds based on genomic analyses and experimental reports, and the genes are summarized in Table 1J. Based on information linking specific precursors to glycan types (Fig. 8D), genetic disruption of a precursor pathway offers an alternative approach to investigate the functional significance of a glycosylation pathway with multiple redundant GT genes.

**Fig. 8.**
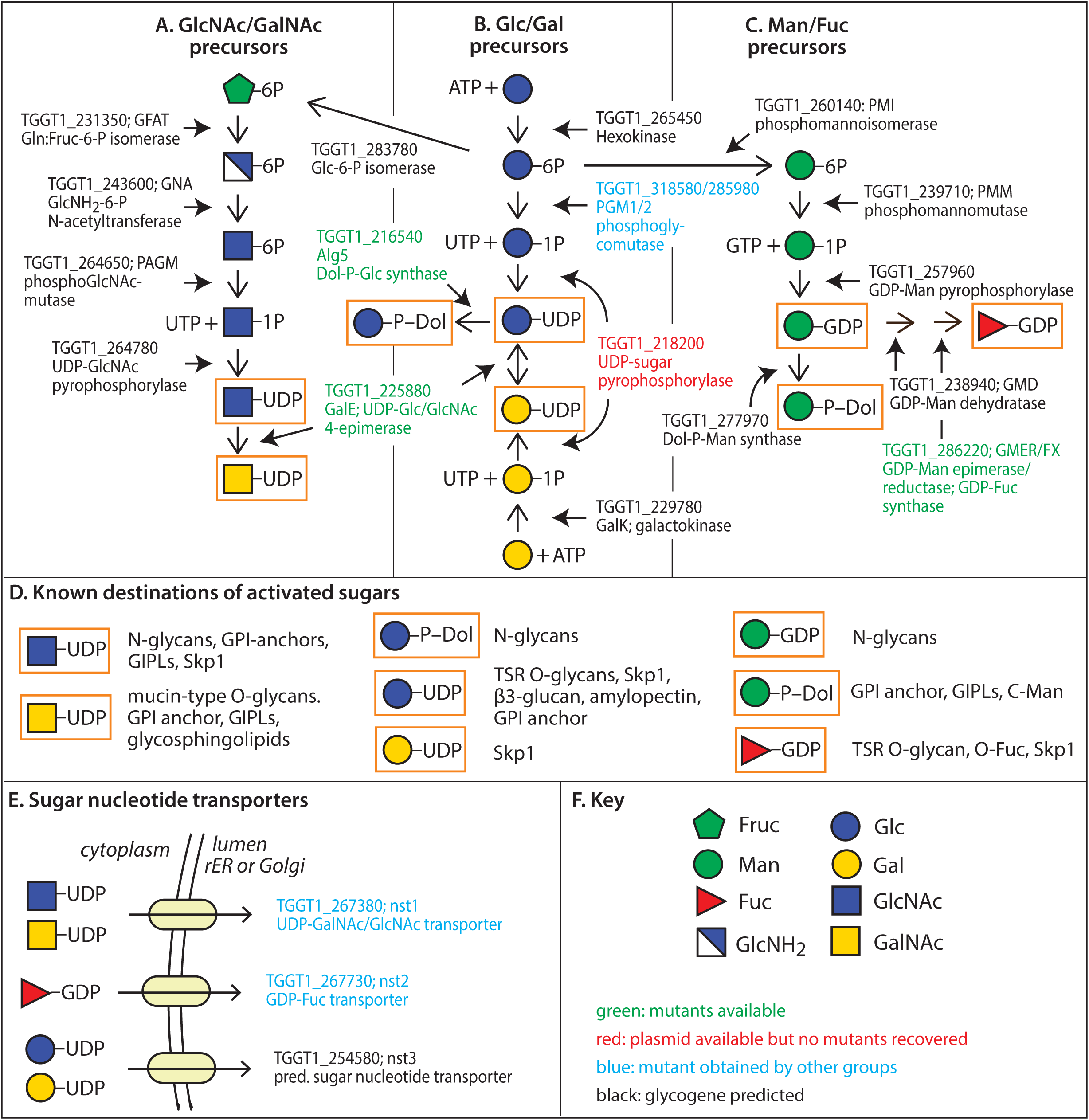
Precursor assembly pathways and transporters. Predicted and validated glycogenes associated with formation and transport of sugar nucleotides, and formation of dolichol-linked monosaccharides, that are utilized as sugar donors by glycosyltransferases. (**A**) UDP-GlcNAc and UDP-GalNAc are derived from Glc via Glc6P. (**B**) UDP-Glc, UDP-Gal, and Dol-P-Glc can derive from either Glc or Gal. (**C**) GDP-Man, Dol-P-Man and GDP-Fuc also ultimately derive from Glc. (**D**) Destination glycans for the precursors boxed in orange in panels A-C. (**E**) Transporters transfer sugar nucleotides from the cytoplasm where they are synthesized into the lumina of vesicles containing the glycosyltransferases. The scramblases that re-orient Dol-P-Glc and Dol-P-Man from the cytoplasmic to the luminal face of the rER are unknown. (**F**) Definition of symbols and color coding of the status of gene analyses.

Current information suggests that all sugars normally derive from Glc or Gal (Fig. 8B). Glc enters the cell via the plasma membrane associated Glc transporter TgGT1 (Zineker et al., 1998), and the α-anomer is converted to Glc-6-PO_4_ (Glc6P) by hexokinase (Saito et al., 2002). Glc6P can be the source of all glycosylation precursors, and is also the C-source for central carbon metabolism via the classical Emden– Meyerhof glycolytic pathway and the pentose phosphate shunt. Alternatively, precursors for glucosylation and galactosylation may derive from the α-anomer of Gal via a salvage pathway (see below), as TgGT1 also transports Gal, Man and GlcNAc (Blume et al., 2009). Finally, glutamine can contribute to sugar precursor assembly via Glc generated by gluconeogenesis (Blume et al., 2015).

The lipid linked *N*-glycan donor contains GlcNAc, Man and Glc, whose precursors derive from each of the three main branches of precursor assembly (Fig. 8A-C). GlcNAc derives from UDP-GlcNAc which is assembled from Glc6P using the classical hexosamine biosynthetic pathway involving 5 enzymes (Fig. 8A). The highly negative fitness score (-2.5 to -5.2) of several of the enzymes implies essentiality of the pathway, consistent with the importance of *N-*glycosylation, so the significance of the less negative fitness score (-1.4) of the middle enzyme GNA is unclear. The ability to radiolabel glycans with tritiated GlcNH_2_ (Azzouz et al., 2006; Azzouz et al., 2002), which presumably requires a transporter and a 6-hexokinase to enter the pathway, is however not explained by current knowledge. Owing to the likely necessity of these enzymes for viability (Cova et al., 2018), UDP-GlcNAc formation was not targeted in this study.

Man derives from GDP-Man, which is formed via three conventional enzymatic transformations from Glc6P (Fig. 8C). The strongly negative fitness scores (-2.7 to -5.0) also suggest essentiality, consistent with the fitness scores for the *N*-glycan ManTs. The very negative fitness score for *pmi* (-5.0) indicates that Glc6P is the only C-source for this pathway. Thus GDP-Man biosynthesis was not targeted in this study. The ability of parasites to be radiolabeled with tritiated Man is consistent with the activities of the Glc transporter and hexokinase to utilize this sugar in addition to Glc.

The GlcNAc and Man residues of *N*-glycans are incorporated while the lipid-linked precursor is oriented toward the cytoplasm and derive from sugar nucleotide donors (Aebi et al., 2010). In contrast, the Glc residues are assembled after the precursor is flipped to the luminal face by a scramblase whose identity is unknown in any eukaryote. As is typical, the 3 αGlcT’s, Alg6, Alg8 and Alg10, utilize Dol-P-Glc as donors. Dol-P-Glc is formed from UDP-Glc by Dol-P-Glc synthase (Alg5), whose -3.4 fitness score suggests importance of this modification, because there is no evidence that other GTs utilize this precursor. *alg5* was successfully disrupted, which resulted in the absence of the Glc modifications and small plaque sizes as expected (see *N*-glycans above).

UDP-Glc is in turn assembled from Glc6P via two traditional enzymatic transformations (Fig. 8B). Conversion to Glc1P is expected to be mediated by a phosphoglucomutase (Imada et al., 2010), but a double-KO of the two candidate genes does not affect viability (Saha et al., 2017). Therefore, this function might also be provided by phospho-N-acetylglucosamine mutase (*pagm*) or phosphomannomutase (*pmm*), as these enzymes are promiscuous in other parasites (Bandini et al., 2012). There is a single UDP-sugar pyrophosphorylase (*usp*) candidate whose score (-0.9) suggests that it also does not contribute much to fitness. Since this varies from the negative fitness score for Dol-P-Glc synthase, the UDP-GlcNAc pyrophosphorylase may be promiscuous as in *Arabidopsis* (Yang et al., 2010) allowing for redundancy. Nevertheless we have been unsuccessful in disrupting this gene using our double CRISPR approach.

Alternatively, Gal may be an alternate source of UDP-Glc via the so-called Isselbacher pathway found in plants and other eukaryotic parasites (Lobo-Rojas et al., 2016). The Glc transporter TgGT1 also transports Gal (Blume et al., 2009) and *Toxoplasma* has predicted galactokinase (GalK; fitness -2.2) and UDP-Glc/UDP-Gal epimerase (GalE; fitness +1.7) enzymes. There is no good candidate for the Gal1P uridyltransferase of the classical Leloir pathway, suggesting that USP, the pyrophosphorylase that forms UDP-Glc from Glc1P, may promiscuously perform this role as observed in a trypanosome and plants (Damerow et al., 2010; Yang and Bar-Peled, 2010). The importance of this pathway was investigated by disrupting *galE*. This deletion resulted in a rather strong plaque defect (Fig. 3) despite a neutral fitness score, consistent with disruption of the Skp1 GalTs. However, GalE is also involved in the formation of UDP-GalNAc from UDP-GlcNAc as demonstrated by effects on mucin-type *O*-glycosylation (Fig. 5G). A selective effect on UDP-Gal formation could be investigated by disrupting GalK, for which a double CRISPR plasmid was generated. Thus *Toxoplasma* may utilize redundancy in its phosphoglycomutases and UDP-sugar pyrophosphorylases, as observed in other eukaryotes, to ensure access to precursors within varied host cells. Alternatively, other genes may yet to be identified, or a critical intermediate might derive directly from the host cell.

Assembly of the GPI anchor backbone, and *C*-Man of TSR domains, depend on Dol-P-Man. The Dol-P-Man synthase gene was not targeted owing to its very negative fitness score (-4.3) and the likely essentiality of the GPI anchor glycan (Fig. 6).

Fuc is known to be selectively applied to TSR domains in the secretory pathway, and to Skp1 and over 60 other proteins in the nucleus and cytoplasm. *Toxoplasma* lacks good candidate genes for the salvage pathway for GDP-Fuc assembly but does possess the two genes for the so-called conversion pathway. The second enzyme gene, the epimerase (or GDP-Fuc synthase), was modified by an insertion at near the 5’- end of its coding region yielded a modest small plaque phenotype (Fig. 3). However, the Glc-Fuc-structure was expressed normally, emphasizing the value of the double CRISPR method and the importance of biochemically confirming consequences of genetic modifications.

Altogether, the metabolic pathways summarized in Fig. 8 that generate activated sugars in *Toxoplasma* are thus far consistent with the gene repertoire and biochemical studies. However, it is still not clear how parasites can be labeled with free tritiated Man and GlcNH_2_ and how tritiated Gal can label GalNAc (Azzouz et al., 2002). Nevertheless, the gene assignments resemble those of the agent for malaria, the related apicomplexan *Plasmodium* (Cova et al., 2015). Two major differences are that i) *Toxoplasma* also possesses enzymes for the utilization of Gal, consistent with the ability to metabolically label the parasite with tritiated Gal (Azzouz et al., 2002), and good candidate genes for GalK and GalE, and ii) *Toxoplasma* has a Dol-P-Glc synthase to glucosylate its *N*-glycans within the rER lumen.

The sugar nucleotides described above are synthesized in the cytoplasm. Their access to GTs within the secretory pathway is afforded by specific transmembrane sugar nucleotide transporters (Fig. 8E), of which three are predicted in *Toxoplasma* (Table 1K). Nst1 has been examined biochemically and shown genetically to be important for the availability of UDP-GalNAc for *O*-glycosylation and for UDP-GlcNAc (Caffaro et al., 2013). Recent studies demonstrated a role for Nst2 in the transport of GDP-Fuc (Bandini et al., 2018). Nst3, which has a very negative fitness score (-4.9) has yet to be studied and may contribute to the transport of UDP-Gal and/or UDP-Glc, but biochemical studies are required to evaluate this speculation.

### CONCLUSIONS

The *Toxoplasma* genome encodes at least 37 predicted GTs, and one phospho-GT, that modify a variety of proteins and lipids or generate polysaccharides. Twenty-six appear to be associated with the secretory pathway and 11 with cytoplasmic glycosylation. An additional 22 genes appear to be involved in generation of glycosylation precursors and their transport into the secretory pathway. This enzyme gene enumeration creates a new opportunity to explore gene-glycan-cell function relationships.

The range and abundance of *Toxoplasma* glycan classes, as profiled by mass spectrometry and including findings from radiolabeling, mAb and lectin studies, is more than initially appreciated but relatively limited compared to other protozoans such as trypanosomatid parasites and the amoebozoan *Dictyostelium* (Rodrigues et al., 2015; Sucgang et al., 2011; Feasley et al., 2015; Schiller et al., 2012). Furthermore, there is notable lack of known anionic modifications of *Toxoplasma* glycans that are prominent elsewhere in the form of anionic sugars or substitutions with PO_4_ or SO_4_. These differences may be a consequence of the homogenous environment experienced by this parasite, because when it is not residing within the parasitophorous vacuole of its host cell, it is enclosed within a protective cell wall.

*N*-glycosylation is remarkably simple, with incomplete Glc-trimming and little evidence for further remodeling of the novel Man_6_GlcNAc_2_-precursor, consistent with the absence of *N*-glycan mediated quality control mechanisms in the parasite rER (Bushkin et al., 2010; Samuelson and Robbins, 2015). *O*-glycans of the *Toxoplasma* secretory pathway are also simple, consisting mainly (if not only) of GalNAc-GalNAc- and GalNAc- in the mucin class, and Glc-Fuc- and Fuc- in the TSR class. *Toxoplasma* pp αGalNAcTs are more related to animal pp αGalNAcTs (CAZy GT27) than typical protist pp αGlcNAcTs (CAZy GT60) from which they evolved (West et al., 2004), suggesting that they derived from a host by horizontal gene transfer. This interpretation is consistent with the uniquely apicomplexan presence of the TSR glycosylation genes, and the evident absence of more diverse *O*-glycosylation, though the presence of four uncharacterized GT-like genes allows for the possibility of additional complexity at lower abundance or at other stages of the parasite life cycle.

Distinct *O*-glycosylation pathways occur in the parasite nucleus and cytoplasm. Over 60 different nucleocytoplasmic proteins have been shown to be modified by *O*-Fuc (Bandini et al., 2016) in a pattern that is reminiscent of *O*-GlcNAc modifications that occurs on thousands of animal nucleocytoplasmic proteins. This parallel is emphasized by the paralogous relationship, presented here, of their enzyme genes, which is certain to expose new mechanisms of protein regulation as has unfolded from studies on *O*-GlcNAc. The cytoplasm also harbors 5 additional GT activities that assemble a pentasaccharide on a single protein (West and Blader, 2015), Skp1, which contrasts with the role of the *O*-Fuc (Spy) GT to add a single sugar to many proteins. The cytoplasm rather than the apicoplast plastid is the locus of the enzymes that process amylopectin (starch), and additional unknown glycans are suggested by 3 other unknown cytoplasmic GT-like genes.

Altogether, glycosylation has many clear roles in the parasite that remain to be explored. The double-CRISPR approach is designed to facilitate these studies by its applicability to all wild-type strains of *Toxoplasma* (owing to NHEJ DNA repair), but the validated guide DNAs are readily adaptable to use in Ku80Δ strains lacking NHEJ, by using PCR to append 45-bp gene-specific homology arms to the ends of the DHFR amplicon. Use of Ku80Δ strains may limit promiscuous insertions of the DHFR expression cassette in locations other than that targeted by the pair of guide RNAs. All guide sequences, CRISPR/Cas9 plasmids, and disrupted strains as summarized in Tables S1 and S4 are available to the community; individuals may contact C.M.W. It is anticipated that these strains and tools will be useful for others to investigate contributions of glycosylation to their favorite cellular or protein functions.

## SUPPLEMENTAL INFORMATION

Tables S1-S5, and Figures S1-S7.

## AUTHOR CONTRIBUTIONS

HTI and MIS designed and generated the plasmids and most of the gene edited cell lines, and validated the target gene loci. BD and MM created and characterized additional gene edited lines. EGP prepared the samples for glycomic studies, performed independent genomic confirmation, interpreted the MS glycomic data, and correlated the data. MOS performed the MS glycomic studies. GB and JS characterized the spyΔ strain and assisted in glycogene identifications. CMW and LW designed the study. CMW supervised the glycogene predictions and wrote the manuscript, which was edited by EGP, MOS, GB, JS, and approved by all authors.

## ACKNOWLEDGMENTS

We thank Bernard Henrissat (CNRS, Marseilles) and Kazi Rahman (Georgia) for their assistance in glycogene identifications. We are grateful to Drs. Tadakimi Tomita and Louis Weiss for the pp GalNAcT mutants, and to Drs. Ulla Mandel and Henrik Clausen for the gift of anti-Tn mAbs. This work was funded by a grant from the NIH Common Fund to Promote the Glycosciences (5R21 AI123161), and received additional support from NIH R01-037539 and NIH R01-084383.

## Abbreviations

AAL: *Aleuria aurantia* agglutinin or lectin
Dol-P-sugar: a conjugate of Glc or Man to dolichol phosphate
EIC: extracted ion chromatogram
FBS: fetal bovine serum
GPI: glycophosphatidylinositol
GT: glycosyltransferase
HAc: acetic acid
HdH: a disaccharide consisting of a hexose and a deoxy-hexose residue
HFF: human foreskin fibroblasts
hTERT: HFF immortalized with human telomerase reverse transcriptase
H6N2: an oligosaccharide consisting of 6 hexose (Hex) and 2 N-acetylhexosamine (HexNAc) residues
NHEJ: non-homologous end-joining
nLC-MS/MS: nano-liquid chromatography coupled to multi-dimensional mass spectrometry
mAb: monoclonal antibody
pp- αGalNAcT: UDP-GalNAc-dependent polypeptide α-N-acetyl-D-galactosaminyltransferase
Tn antigen: a structure corresponding to αGalNAc-S/T
TSR: thrombospondin type 1 repeats

